# Chiari II brain malformation is secondary to open spina bifida

**DOI:** 10.1101/2025.01.06.631442

**Authors:** Maryam Clark, Timothy J. Edwards, Dawn Savery, Gabriel L. Galea, Nagaraj Samy, Erwin Pauws, Nicoletta Kessaris, Nicholas D.E. Greene, Andrew J. Copp

**Affiliations:** Developmental Biology & Cancer Department, Great Ormond Street Institute of Child Health, University College London, UK; Wolfson Institute for Biomedical Research, University College London, UK; Metro South Health, Queensland, Australia; MRC National Mouse Genetics Network, Congenital Anomalies Cluster, Mary Lyon Centre at MRC Harwell, UK

**Keywords:** neural tube defect, myelomeningocele, mouse, fetus, transgenic, skull

## Abstract

Chiari II brain malformation affects 90% of children with open spina bifida. Hindbrain herniation leads to hydrocephalus, together with higher brain anomalies including cerebral cortical defects implicated in learning disability, which affects 20-25% of children with spina bifida. The causal link between Chiari II and spina bifida has long been debated, and we aimed to determine whether Chiari II arises secondary to spina bifida, rather than as a separate effect of shared genetic or non-genetic factor(s). *Pax3* gene function was conditionally deleted by *Cdx2^cre^* specifically in the lower body of mice, leaving the head genetically intact. Open spina bifida is seen in all *Cdx2^cre/+^; Pax3^fl/fl^*fetuses, together with many features of Chiari II in the wild-type brain and skull. These include: hindbrain herniation, callosal and hippocampal hypogenesis, cortical thinning with neuronal heterotopia, a thickened ventricular zone, and posterior skull defects. Hence, the brain and skull defects of Chiari II arise secondary to open spina bifida, with likely disturbance of neurogenesis and neuronal migration early in gestation. The *Cdx2^cre/+^; Pax3^fl/fl^* mouse provides a model for improved understanding of Chiari II pathogenesis.

## INTRODUCTION

The Chiari II congenital brain malformation affects up to 90% of individuals with open spina bifida (SB; myelomeningocele). The cerebellar vermis, with or without the brainstem, herniates through the foramen magnum into the vertebral canal. This can lead to hindbrain compression that may cause acute bulbar dysfunction with impaired swallow, aspiration, inspiratory stridor and central apnoea. Posterior fossa surgical decompression may be required in such emergency cases. More chronically, Chiari II is strongly associated with subsequent development of hydrocephalus, which often requires placement of a cerebrospinal fluid (CSF) shunt, and periodic revision surgery (2003; Stevenson, 2004; Tubbs and Oakes, 2013).

Chiari II is not confined to hindbrain herniation but is a global brain syndrome with, in addition to hydrocephalus, medullary kinking and malformations of the ‘higher’ (supratentorial) brain including an enlarged massa intermedia of the thalamus, small third ventricle, tectal beaking, dysgenesis of the corpus callosum and defects of cerebral cortical structure (Copp et al., 2015). The severity of these brain defects correlates with neurocognitive deficit which affects 20-25% of children with SB, and can seriously affect the lives and independence of people with SB (Rofail et al., 2013; Schneider et al., 2021).

The strong association of Chiari II with SB has prompted much speculation in the neurosurgical and other literature on the causal relationship between the two conditions. In principle, they may share a common cause: for example, a genetic factor that affects both brain and spine directly, so that SB and Chiari II occur frequently together but are largely independent of each other developmentally. Alternatively, Chiari II could be a secondary consequence of SB, and this idea has been elaborated in most detail within the ‘unified’ hypothesis of McLone and Knepper (1989). The persistent leakage of cerebrospinal fluid (CSF) from the SB lesion is seen as preventing establishment of a sealed ventricular system within the brain, with reduced hydrostatic pressure causing failure of the developing hindbrain to expand normally. Skull tissues are ‘induced’ by the adjacent brain structure during development (Richtsmeier and Flaherty, 2013), so the deflated hindbrain is seen as inducing an abnormally small posterior cranial fossa. This cannot contain the cerebellum and brain stem which herniate through the foramen magnum as fetal development proceeds.

In recent years, the widespread introduction of surgery for SB in the fetal period has led to the finding that the hindbrain herniation of Chiari II can be diminished, in the short-term following closure of the SB lesion, with a significant reduction in the need for CSF shunting (Adzick et al., 2011; Joyeux et al., 2018). However, postnatal MRI assessment of children after fetal SB surgery shows that, while hindbrain herniation is reduced, the constellation of higher brain defects is still present in almost all cases (Calle et al., 2020). Hence, the supratentorial aspects of the Chiari II malformation are not ameliorated by fetal surgery in the second trimester, arguing for an earlier developmental origin of the brain defects.

Here, we tested the hypothesis that Chiari II results secondarily from the occurrence of SB. Mouse fetuses that completely lack function of the *Pax3* gene develop SB and, in a preliminary study, we identified co-existing hindbrain herniation by fetal MRI (Norris et al., 2015). In the present study, we generated embryos and fetuses lacking *Pax3* only in the lower body, using *Cdx2^cre^* to recombine a floxed *Pax3* allele in the trunk region. These embryos develop SB at full penetrance, and we demonstrate that both hindbrain herniation and higher brain defects are present in the genetically wild-type heads of these embryos and fetuses. This study shows experimentally for the first time that Chiari II develops secondary to SB, and establishes the *Cdx2^cre^/Pax3^fl^*mouse as a model system in which to study the full constellation of developmental brain anomalies in individuals with SB.

## RESULTS

Chiari II co-exists with most cases of human SB and, in mice, hindbrain herniation has been detected in several SB models involving genetic (Briner and Moellenberndt, 1997; Miro et al., 2009; Norris et al., 2015) or teratogenic (Alles and Sulik, 1992; Briner and Lieske, 1995) influences. However, in none of these animal studies was it possible to determine whether the features of Chiari II were direct effects of the causative gene/teratogen on the brain or, alternatively, whether they arose secondary to the SB. To resolve this long-standing question, we made use of Cre-loxP recombination technology to generate a *Cdx2^cre/+^;Pax3^fl/fl^*mouse. This is a model designed to selectively remove PAX3 expression from the caudal half of the embryo, while leaving the head wild-type (Figure 1A). We reasoned that the presence or absence of Chiari II defects in the brain and skull of *Cdx2^cre/+^;Pax3^fl/fl^* mice with SB would resolve the possible cause-and-effect relationship between SB and Chiari II.

**Figure 1:**
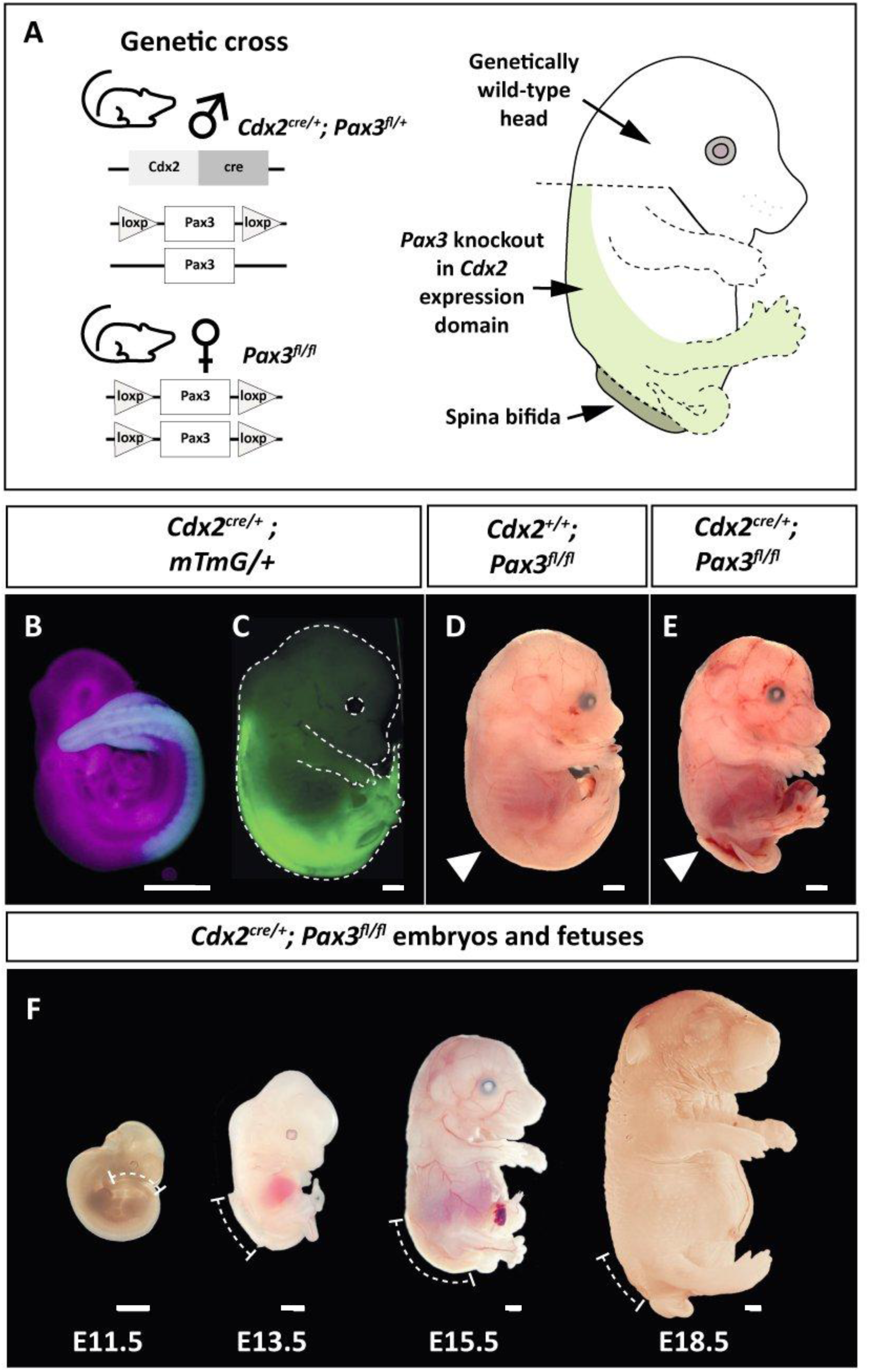
Generation of the *Cdx2^cre^;Pax3^fl/fl^* mouse mousemodel. (**A**) Genetic cross used to generate mice with SB. *Cdx2^cre^* drives *Pax3* knockout in the body only (light green highlight), whereas the head remains wild-type. See Table 1 for offspring genotypes. (**B, C**) *Cdx2^cre^* driving *mTmG* reporter expression in the body at E10.5 (B, cyan) and E15.5 (C, green). Magenta in (B) indicates region of no *Cdx2_cre_*-mediated recombination. See also Supplementary Figure 1. (**D, E**) Representative images of E15.5 control (*Cdx2^+/+^*; *Pax3^fl/fl^)* and mutant (*Cdx2^cre^*^/+^*; Pax3^flf^*) fetuses. White arrowheads indicate the location of the open SB lesion (in E). See also Supplementary Figure 2. (**F**) Phenotypic developmental timeline of *Cdx2^cre/+^;Pax3^flfl^* embryos and fetuses, with dotted lines indicating extent of open SB lesions. All scale bars: 1.0 mm.

**Table 1.** The four genotypes resulting from the matings between *Cdx2^cre/+^; Pax3^fl/+^* males and *Pax3^fl/fl^* females, with embryo numbers and phenotypes from 10 litters (n = 78 embryos).

|  | No. of embryos | % all embryos * | No. normal | No. SB |
| --- | --- | --- | --- | --- |
| *Cdx2<sup>cre/+</sup>*; *Pax3<sup>fl/fl</sup>* | 15 | 19.2 | 0 | 15 |
| *Cdx2<sup>cre/+</sup>*; *Pax3<sup>fl/+</sup>* | 22 | 28.2 | 22 | 0 |
| *Cdx2<sup>+/+</sup>*; *Pax3<sup>fl/fl</sup>* | 12 | 15.4 | 12 | 0 |
| *Cdx2<sup>+/+</sup>*; *Pax3<sup>fl/+</sup>* | 29 | 37.2 | 29 | 0 |

\* Statistical analysis: Observed genotype ratios do not differ significantly from Mendelian expectation of 25% for each genotype (Chi-square 4 x 2 test; p = 0.171).

### *Cdx2*^cre^ recombines *Pax3* solely in the lower body

Use of *mTmG* and Rosa26-eYFP reporter lines confirmed the previous finding (Coutaud and Pilon, 2013) that *Cdx2^cre^*-mediated recombination is confined to the lower embryonic and fetal body. We detected a rostral recombination limit at upper trunk (forelimb) level at E10.5 (Figure 1B), E15.5 (Figure 1C) and E18.5 (Supplementary Figure 1). Open SB occurred in *Cdx^cre/+^;Pax3^fl/fl^*embryos and fetuses (Figure 1D-F; Supplementary Figure 2), consistent with the SB observed in *splotch* mutant mice that constitutively lack PAX3 (Greene et al., 2009). To determine whether, as predicted, PAX3 persists in the heads of *Cdx^cre/+^;Pax3^fl/fl^*embryos, we performed anti-PAX3 immunohistochemistry on sagittal sections of E10.5 embryos. Similar intensities of PAX3 expression were detected in the heads of control (non-Cre) and *Cdx^cre/+^;Pax3^fl/fl^*embryos, and also in the lower body of controls (Figure 2A-C). Strikingly, however, there was no PAX3 signal in the lower body of mutants, demonstrating the loss of PAX3 expression in association with open SB in the mouse model (Figure 2D). These findings confirm that PAX3 expression has been selectively removed from the trunk region of *Cdx2^cre/+^;Pax3^fl/fl^* embryos, but remains intact in the head.

**Figure 2:**
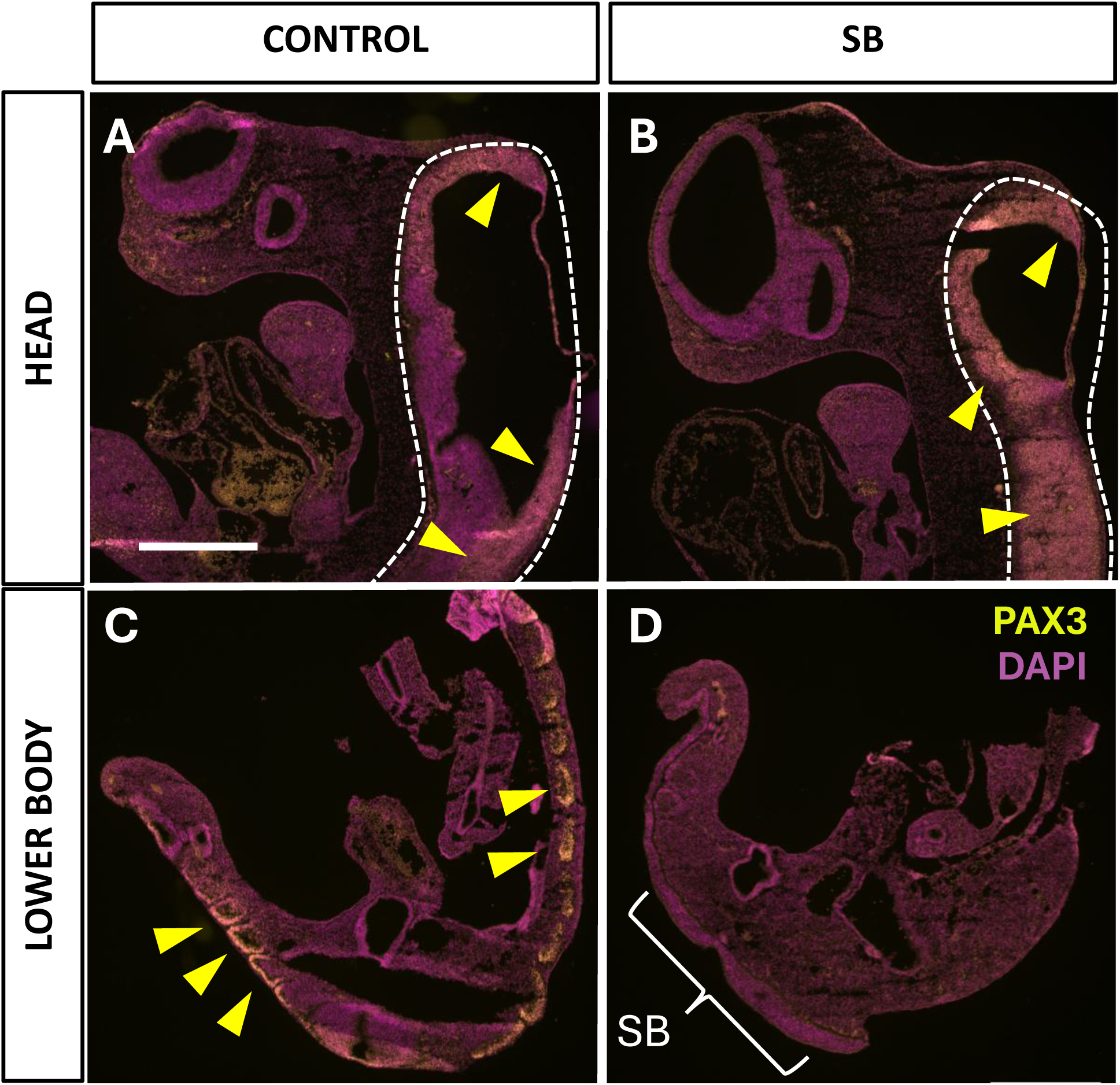
PAX3 expression in E10.5 control and *Cdx2^cre/+^;Pax3^fl/fl^*embryos. Sagittal cryosections of control (n = 3) and SB (n = 3) embryos, immunostained for PAX3 (yellow) and nuclear-stained with DAPI (magenta). (**A, B**) Head sections: PAX3 expression is present in the neural tube (hindbrain, outlined by dashed lines) of both control and SB (yellow arrowheads). (**C, D**) Lower body: PAX3 expression is present in the dorsal neural tube and dermomyotomes of the control spinal region (yellow arrowheads in C), but is not detectable in the SB embryo (D), consistent with knockout of *PAX3* in the lower body. Scale bar for all parts: 0.5 mm.

It was important to examine the efficiency of *Cdx^cre^*-mediated *Pax3* removal in inducing SB, particularly as different *splotch* alleles (e.g. *Pax3^Sp^*, *Pax3^Sp2H^*, *Pax3^d^*) vary in their penetrance of SB (Greene et al., 2009). Among 10 sampled litters, from *Cdx^cre/+^;Pax3^fl/+^* x *Pax3^fl/fl^* matings, we identified 15 *Cdx^cre/+^;Pax3^fl/fl^* (hereafter ‘mutant’) embryos and fetuses, all of which exhibited SB (Table 1). Hence, *Cdx^cre/+^;Pax3^fl/fl^* genotype produces fully penetrant SB.

### Hindbrain herniation in the *Cdx2^cre/+^;Pax3^fl/fl^* mouse

In children with SB, cerebellar herniation and a small posterior fossa with enlarged foramen magnum are typical signs of the Chiari II malformation. To explore their occurrence in the mouse model, we carried out histology and performed micro-computed tomography (microCT) for both hard and soft tissues. In haematoxylin and eosin (H&E)-stained midsagittal sections of E18.5 brains, the distance between the lowest point of the cerebellum and a line drawn from the base of the skull to the posterior part of the parietal bone (hereafter referred to as the ‘skull boundary’) was compared between control and mutant fetuses (Figure 3A, B; Supplementary Figure 3). In addition, the dorso-ventral length of the skull boundary length was measured as equivalent to the foramen magnum at this stage. SB mutants exhibit a highly significant cerebellar herniation compared with control fetuses whose cerebellum is confined within the skull (Figure 3C). In contrast, there was no significant difference between control and SB fetuses in length of the skull boundary (Figure 3D). Parietal bone length, normalised to head length, did not differ between control and SB fetuses (Supplementary Figure 3). Hence, hindbrain herniation is present in the *Cdx2^cre/+^;Pax3^fl/fl^* mouse model, as in human Chiari II.

**Figure 3:**
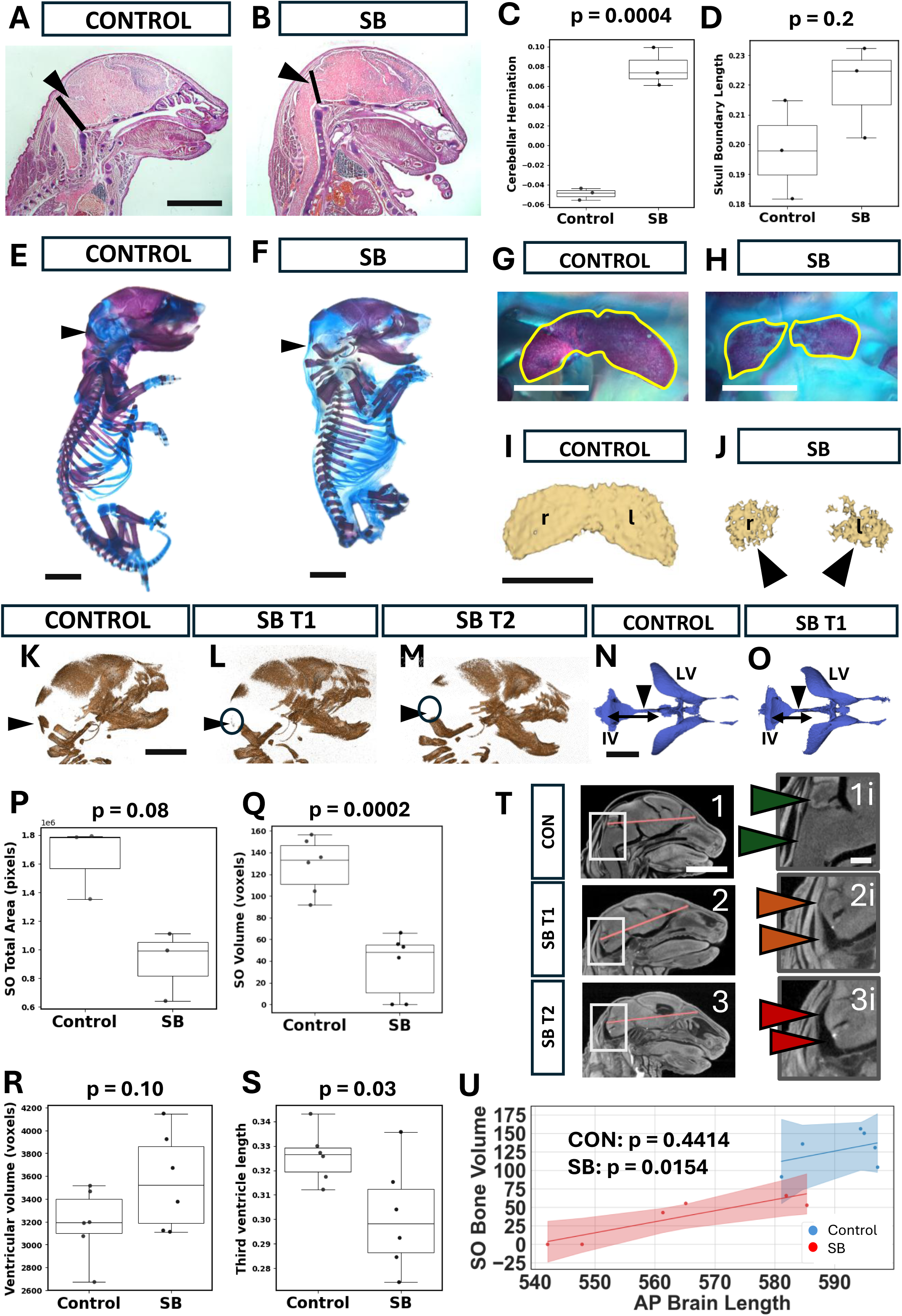
Hindbrain herniation and supra-occipital (SO) bone formation. (**A,B**) H&E-stained head sagittal sections at E18.5. See also Supplementary Figure 3 for higher magnification views of skull and brain. A line was drawn from the caudal edge of the parietal bone to skull base (purple tissue, black line), and taken as the boundary between skull and spinal region at this stage. Distances between the cerebellum (black arrowheads) and the line were measured: below/caudal to the line = positive values; above/rostral = negative values. The cerebellum in SB is located below the line (B), indicating a herniation. Scale bar: 2 mm. (**C, D**) Boxplots showing comparisons between control and SB (images as in A, B) for cerebellar position relative to the boundary (C), and length of the boundary line between skull and spinal region (D). Values were normalised to head A-P length. Cerebellar position is significantly more caudal (positive values) in SB than control (C; Welch’s t-test: p = 0.0004) whereas boundary length does not differ between control and SB (D; Mann-Whitney U Test: p = 0.1763). (**E, F**) Right-sided views of alizarin red (bone) and alcian blue (cartilage) stained skeletal tissues in control and SB at E18.5. Black arrowheads indicate the region of the SO bone. Scale bars: 2 mm. (**G-J**) Dorsal views of SO bone in control and SB skull preparations (G, H) and after microCT 3D-reconstruction (I, J). Alizarin red-stained bone is outlined in yellow (G, H). Arrowheads: rudimentary SO bones in SB as visualised by microCT. r: right; l: left. Scale bars: 0.5 mm. (**K-M**) MicroCT 3D-reconstructions of the cranium in control, SB T1 (SO present) and SB T2 (SO absent) at E18.5. Black arrowheads: region where the SO bone is usually found, although markedly reduced or absent in SB (black circles). Scale bar: 2 mm. (**N, O**) MicroCT 3D-reconstructions of brain ventricular system in control and SB T1 at E18.5. Black arrowheads point to 3^rd^ ventricles, with double-headed arrows indicating the 4^th^ (IV) to 3 ventricular length of these regions, which is reduced in SB. LV: lateral ventricles. Scale bar: 2 mm. (**P-S**) Boxplots showing comparisons (E18.5) of total SO stained area from skeletal preparations (P; n = 3), SO volumes from microCT skeletal preparations (Q; n = 6; normalised to head length), and ventricular volumes (R) and 3 to 4 ventricle lengths (S) from microCT soft tissue 3D-reconstructions (n = 6; normalised to brain length). SO stained area (P) shows a trend towards reduction in SB (Mann-Whitney U test; p = 0.08). SO volume (Q) is significantly reduced in SB (Welch’s t-test; p = 0.0002), as is 3 to 4 ventricle length (S) (Welch’s t-test; p = 0.03), whereas ventricular volume (R) does not differ significantly between control and SB (Welch’s t-test; p = 0.1). (**T**) MicroCT sagittal cranial soft tissue images showing morphological variation in contact between cerebellum/hindbrain and overlying SO bone region. Boxed areas in the left-side images (1-3) are shown at higher magnification in the right-side images (1i-3i). Arrow colours indicate increasing distance (green, amber, red) between cerebellum/hindbrain and SO bone region. Control head shows extensive contact (1i), SB T1 shows limited contact (2i), and SB T2 shows no contact (3i). Scale bars: 2 mm in 1-3; 0.5 mm in 1i-3i. (**U**) Relationship between A-P brain length and SO bone volume in control and SB heads (n = 6 each) at E18.5. A linear correlation exists in SB heads (R = 0.89; p = 0.0154), but not in control (R = 0.18; p = 0.4414), with SB showing significantly shorter brains than controls (Welch’s t-test; p = 0.0082), as well as smaller SO bone volumes (Q).

### Posterior skull defects in the *Cdx2^cre/+^;Pax3^fl/fl^* mouse

In mice, the posterior skull fossa is composed of multiple bones that eventually fuse, including the supraoccipital (SO), which sits directly behind the cerebellum. To visualise SO size and structure, we carried out alcian blue and alizarin red staining of skeletal preparations from control and SB fetuses at E18.5 (Figure 3E, F). The surface area of the SO was measured on photographic images taken from a dorsal/posterior direction. In controls, the SO was a bilobed structure that straddled the midline (Figure 3G), whereas in all three mutants the SO was underdeveloped and existed as two separate bony structures, either side of the midline (Figure 3H). A comparable control-SB difference was found in microCT hard tissue scan images (Figure 3I, J). Quantitative analysis revealed a trend towards a smaller SO area in SB skeletal preparations (Figure 3P) and SO bone volume measured from microCT scans was significantly smaller in SB than control skulls (Figure 3Q).

Close inspection of skull microCT scans revealed similar degrees of overall ossification in control and SB skulls (Figure 3K-M). Moreover, two degrees of SO dysgenesis could be detected in fetuses with SB: partial (SB T1; Figure 3L) and complete (SB T2; Figure 3M). To determine possible reasons for this finding, we asked whether lower pressure, resulting from smaller ventricles in the mutants, might induce smaller SO bones. However, when normalised to A-P brain length, overall ventricular volume did not vary significantly between control and mutant brains (Figure 3R), with no correlation between SO volume and ventricular volume in individual fetuses.

An alternative possibility was that variations in direct pressure of the brain on bone-forming cranial mesenchyme might account for the differing degrees of SO dysgenesis. When visually compared in individual fetal microCTs, we could identify little intervening space between the cerebellum/brainstem and skull in controls, with contact between posterior skull and both cerebellum and brainstem (Figure 3T-1,1i). In SB T1, where the SO was reduced in size, there was increased space, with only the cerebellum in direct contact with the skull (Figure 3T-2, 2i). In SB T2, that completed lacked the SO, there was essentially no hindbrain-skull contact at all (Figure 3T-3, 3i). Hence, it seems possible that failure of the hindbrain to contact the developing posterior skull results in the SO dysgenesis we observe in SB heads.

Although overall ventricular volume did not differ between control and SB brains, we found variation in length of the third ventricle in the three-dimensional (3D) microCT reconstructions (Figure 3N, O) with a significant length reduction in SB mutants (Figure 3S). This suggested a possible difference in A-P brain length, and this was compared with SO bone volume using an ordinary least squares regression model, which revealed a strong linear relationship between brain length and SO bone volume in SB heads (Figure 3U: red line and points). Controls exhibited larger A-P brain lengths than SB fetuses, with less variation between individuals (Figure 3U: blue line and points). We conclude that two major signs of Chiari II, herniated hindbrain and diminished posterior skull development, are present in *Cdx2^cre/+^;Pax3^fl/fl^* fetuses, validating this mouse as a model of human Chiari II.

### Soft tissue microCT analysis at E18.5

Having identified a diminished A-P brain and skull axis in mutants, we sought to further understand the structural anomalies associated with SB using soft tissue microCT. This method yielded sagittal images of E18.5 fetuses which revealed both large and small open SB lesions (Figure 4B, C), compared with the closed spine of normal controls (Figure 4A). Interestingly, all six fetuses with SB displayed signs of a full bladder (asterisks in Figure 4B, C), which was not seen in controls. Hence, bladder dysfunction may be a feature of mouse SB, as in affected children. We asked whether SO bone volume (Figure 3K-M) may be related to SB lesion length, and performed linear regression analysis which revealed a trend towards small SO volumes in fetuses with large SB lesions (Figure 4D). This possible relationship may indicate an effect of SB severity on skull morphogenesis in Chiari II.

**Figure 4:**
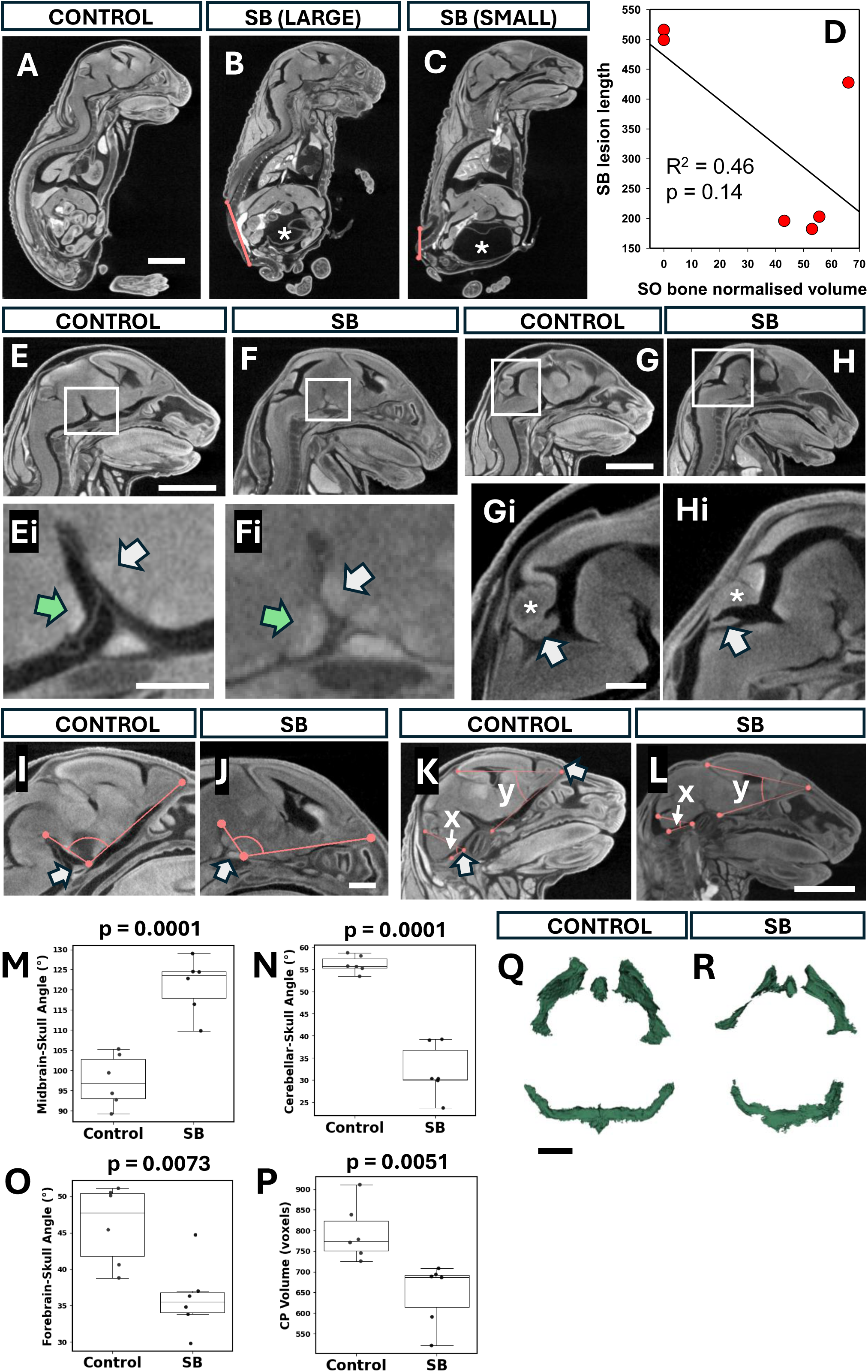
Global signs of brain displacement in SB. (**A-C)** Sagittal images from soft tissue microCT at E18.5 of control (A) and SB fetuses (B, C). Red lines show the size of the SB lesion, which varied from large (B) to small (C). White asterisks indicate a full bladder which was present in all SB fetuses, but not in controls (n = 6 of each). Scale bar: 2 mm. (**D**) Relationship between SB lesion length and SO bone normalised volume. SB fetuses with larger SB lesions tend to have smaller SO bones, although the regression line does not reach statistical significance (p = 0.014) due to the limited sample size (n = 6). (**E-Hi**) Sagittal images of control (E, G) and SB heads (F, H). Boxed areas in (E, F) are shown at higher magnification in (Ei, Fi) revealing significant overlap of pons (green arrows) and hypothalamus (white arrows) in SB (Fi) but not in control (Ei). Magnifications of boxed areas in (G, H) show a flattened cerebellum (asterisk) and choroid plexus (arrow) in SB (Hi) compared with control (Gi). Scale bars: 2 mm in E-H; 0.5 mm in Ei, Fi; 0.5 mm in Gi, Hi. (**I, J**) Sagittal images showing angle between midbrain and plane of forebrain base. SB shows midbrain displacement dorsally, as indicated by larger angle in (J) compared with control (I). White arrows: skull base landmark used for angle normalisation. Scale bar: 0.5 mm in I, J. (**K, L**) Sagittal images to illustrate forebrain “collapse” (reduction in y angle) and cerebellar herniation (reduction in x angle). Both angles are smaller in SB (L) compared with control (K). White arrow: skull base landmark used for angle normalisation. See also Supplementary Figure 5 for higher magnification views of x angle measurements. Scale bar: 2 mm in K, L. (**M-O**) Box plots showing angle data from images in (I-L). Midbrain-skull angle (M: from I, J) is significantly greater in SB than control (Welch’s t-test; p = 0.0001). Cerebellar-skull base angle (N: angle x in K, L) is significantly smaller in SB than control (Welch’s t-test; p = 0.0001), as is forebrain-skull base angle (O: angle y in K, L; Welch’s t-test; p = 0.0073). (**P-R**) Segmented and 3D-reconstructed choroid plexus (CP) from microCT scans: dorsal views with anterior upwards and posterior downwards (Q, R). Quantification (P) shows significant decrease in CP volume in SB (Mann-Whitney U test; p = 0.0051). Scale bar: 1 mm in Q, R.

Chiari II in humans is frequently associated with hydromyelia (La Marca et al., 1997; McLone and Dias, 2003), in which there is enlargement of the spinal cord central canal. We measured central canal width, normalised to spinal cord width, at ten progressively more caudal levels from cervical to thoracic. SB fetuses show an approximate doubling of central canal width over this distance, whereas controls exhibit significantly less change (Supplementary Figure 4). Hence, the mouse SB model displays hydromyelia which could be the direct result of Pax3 deletion, which extends throughout the thoracic level, although persistent CSF leakage through the open SB lesion may also contribute to the increased patency of the central canal.

### Global brain posterior displacement in SB mutants

To address possible anomalies along the brain’s A-P axis, we examined mid-sagittal microCT images and noted an abnormal placement of the pons and hypothalamus (Figure 4E, F). In controls, these structures were clearly separated, whereas in SB mutants, the hypothalamus was compressed against the dorsal element of the pons (Figure 4Ei, Fi). Further, we identified a flattened cerebellum and choroid plexus of the fourth ventricle (Figure 4G-Hi). To quantify these, we measured three angles: midbrain-skull (Figure 4I, J), cerebellum-skull (Figure 4K, L; Supplementary Figure 5; angle x) and forebrain-skull (Figure 4K, L; angle y). Each of these angles was significantly different between control and mutant (Figure 4M-O), suggesting a global posteriorly-directed displacement of the dorsal brain in fetuses with SB.

As the dorsoventral axis of the choroid plexus in the fourth ventricle appeared shortened, we measured total choroid plexus volume in control and mutant fetuses. Morphologically, this appeared smaller in SB mutants (Figure 4Q, R) and this was confirmed when volumes were normalised to A-P brain length (Figure 4P). Together, these findings suggest that SB brains undergo a degree of posteriorly-directed distortion, with reduction in choroid plexus volume.

### Forebrain defects in SB fetuses at E15.5

The immunohistochemical analysis at E10.5 revealed no obvious structural brain alterations in SB mutants versus control embryos (Figure 2A,B). Next, we examined forebrain structure at E15.5, when 5 of the 6 cortical layers have formed, the majority of radial migration has occurred and tangentially migrating neurons have begun entering the neocortex. Coronal sections through control and SB brains were immunostained for CTIP2 and TBR1, markers of cortical layers 5 and 6, with DAPI counterstaining. Whole head sections show a marked dorsoventral ‘compression’ of the SB brain (double arrowheads in Figure 5D-F) compared with controls (Figure 5A-C). At higher magnification, we detected an apparently smaller amygdala region in SB brains (Figure 5G, H), while the lateral ventricles also appeared smaller in SB brains (Figure 5I, J). At the dorsal midline, the TBR1+ cortical plate adjacent to the interhemispheric fissure curves medially in control brains, consistent with onset of callosal fibre crossing (Figure 5K), whereas the interhemispheric TBR1+ cortical plate in SB brains is thickened and appears to diverge from the midline (Figure 5L), suggestive of callosal hypogenesis.

**Figure 5:**
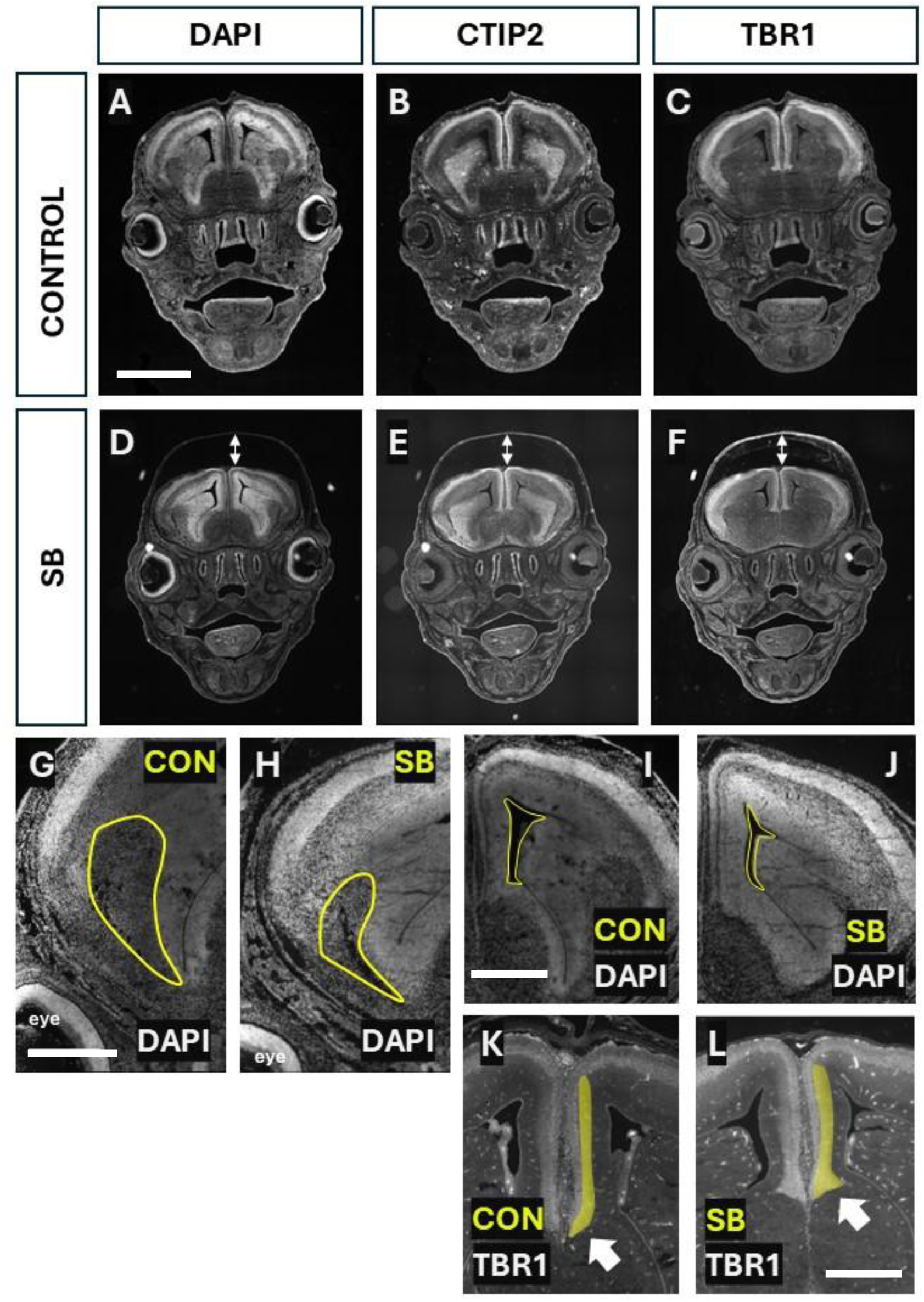
Forebrain defects in SB brains at E15.5. (**A-F**) Coronal sections of control and SB brains, immunostained for CTIP2 and TBR1 with DAPI counterstain. Note the dorsoventrally ‘compressed’ brain in SB (double-headed arrows in D-F), which is absent from control brains (A-C). Scale bar: 2 mm. (**G-L**) Coronal sections stained with DAPI and TBR1 (K, L) showing abnormalities in SB brains compared with control. The developing amygdala (G, H) and the lateral ventricles (I, J) appear smaller in SB (yellow outlines). In the dorsal midline region of control brains, the TBR1+ cortical plate adjacent to the interhemispheric fissure (yellow highlight) converges on the midline (arrow in K), consistent with callosal fibre crossing. In contrast, it is thickened, dorso-ventrally shortened and appears divergent from the midline in SB (arrow in L), suggestive of callosal hypogenesis. Scale bars: 0.75 mm in G, H; 0.5 mm in I, J; 0.5 mm in K, L.

### Differential thickness of cortical plate and ventricular zone in SB

To extend the investigation of brain anomalies associated with mouse SB, we immunostained E18.5 coronal brain sections for the cortical markers, BRN2 and CTIP2 (Figure 6A, B). Comparison of cortical layering revealed a broadly similar pattern in control and SB sections, but with several variations (Figure 6C). The SB cortex appeared thinner overall than control, whereas the ventricular zone (VZ) occupied a larger total proportion of the SB cortical plate than in control. To quantify these differences, we measured total cortical plate and VZ thickness in DAPI-stained brain sections (Figure 6D). Normalising these values to total brain length (smaller in SB: Figure 3U) showed that SB brains have a significantly thinner cortical plate and thicker VZ region than controls, with no differences between right and left sides (Figure 6E, F). We also measured the thickness of layers I-IV, V and VI, after normalisation to total cortical thickness, and identified a significant difference between layers (2-way ANOVA: p < 0.001) but no difference between control and SB (p = 0.132). Together, these results suggest a potential defect in cortical neurogenesis and/or neuronal migration in SB brains.

**Figure 6:**
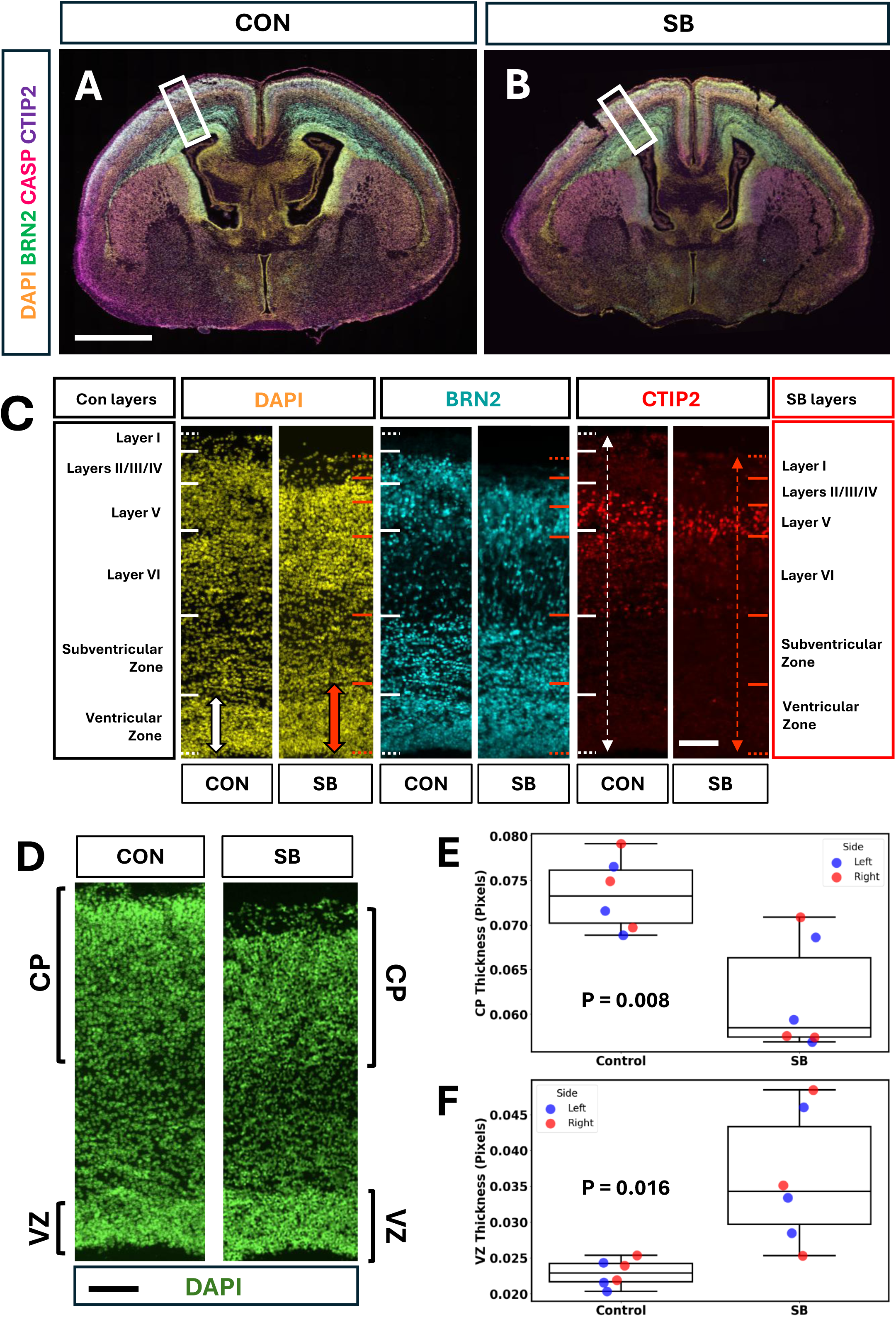
Cortical plate and ventricular zone thickness anomalies in SB at E18.5. (**A, B**) Representative coronal sections of control and SB brains at forebrain level, immunostained for BRN2, CASP and CTIP2, with DAPI counterstain. Scale bar: 1 mm. (**C**) Comparison of cortical layering (inside/apical = bottom; outside/basal = top) in control and SB brain sections (n = 3 per group), immunostained for BRN2 and CTIP2, with DAPI counterstain. White solid lines indicate layer boundaries in control sections, and red solid lines show comparable boundaries in SB sections. Dotted lines mark the inner and outer limits of the forebrain, showing a thinner telencephalic wall in SB than in controls (compare white and red dashed arrows in CTIP2 images). In contrast, the ventricular zone is thicker in SB than control sections (compare filled white and red arrows in DAPI images). Scale bar: 0.05 mm. (**D**) Comparison of relative cortical plate and VZ thickness in control and SB sections with DAPI staining (D). Brackets indicate regions, based on nuclear density, considered as cortical plate and VZ for quantification. Scale bar: 0.05 mm. (**E, F**) Cortical and VZ thickness measurements from left and right sides of n = 3 brains each for control and SB, with normalisation to total brain length. Two-Way ANOVA shows that cortical plate thickness (E) is significantly greater in control than SB (p = 0.008) whereas left (green symbols) and right (red symbols) do not differ significantly (p = 0.71). In contrast, VZ thickness (F) is significantly greater in SB brains (p = 0.016) with no significant difference between left and right sides (p = 0.83).

### Neuronal heterotopia and diminished neuronal number in SB

To test for a neurogenic and/or migratory defect of cells colonising the cerebral cortex, we looked for two signs: a mixing/heterotopia of cortical layers and a decrease in cell number. To test for cortical mixing, we identified a region of interest in the E18.5 cortices of control and SB fetuses (Figure 7A, B) where BRN2+ and CTIP2+ populations appeared more overlapping in SB than in controls (Figure 7C-F). Overlap of these immuno-positive areas was evaluated using a support vector machine-learning algorithm on individual cell coordinates (see Materials and Methods). Figure 7G, H is a visual representation of control and SB algorithmic learning and recall performance, while Figure 7I shows the F1 scores of control and SB samples. F1 scores closer to 1 indicate better predictive accuracy of the algorithm in drawing a ‘perfect’ boundary between the two cell populations, whereas F1 scores close to 0 indicate extensive overlap, with no clear boundary. SB brains performed significantly closer to zero than controls (Figure 7I). Hence, SB brains exhibit diminished segregation of cortical layer populations, indicative of cortical heterotopia. Further, manual cell counts of the BRN2+ and CTIP2+ populations revealed a significant decrease in BRN2+ cells in SB brains (Figure 7J), whereas CTIP2+ cell number did not differ significantly (Figure 7K). Together, these results suggest a neurogenic and/or migratory defect in SB cortical development.

**Figure 7:**
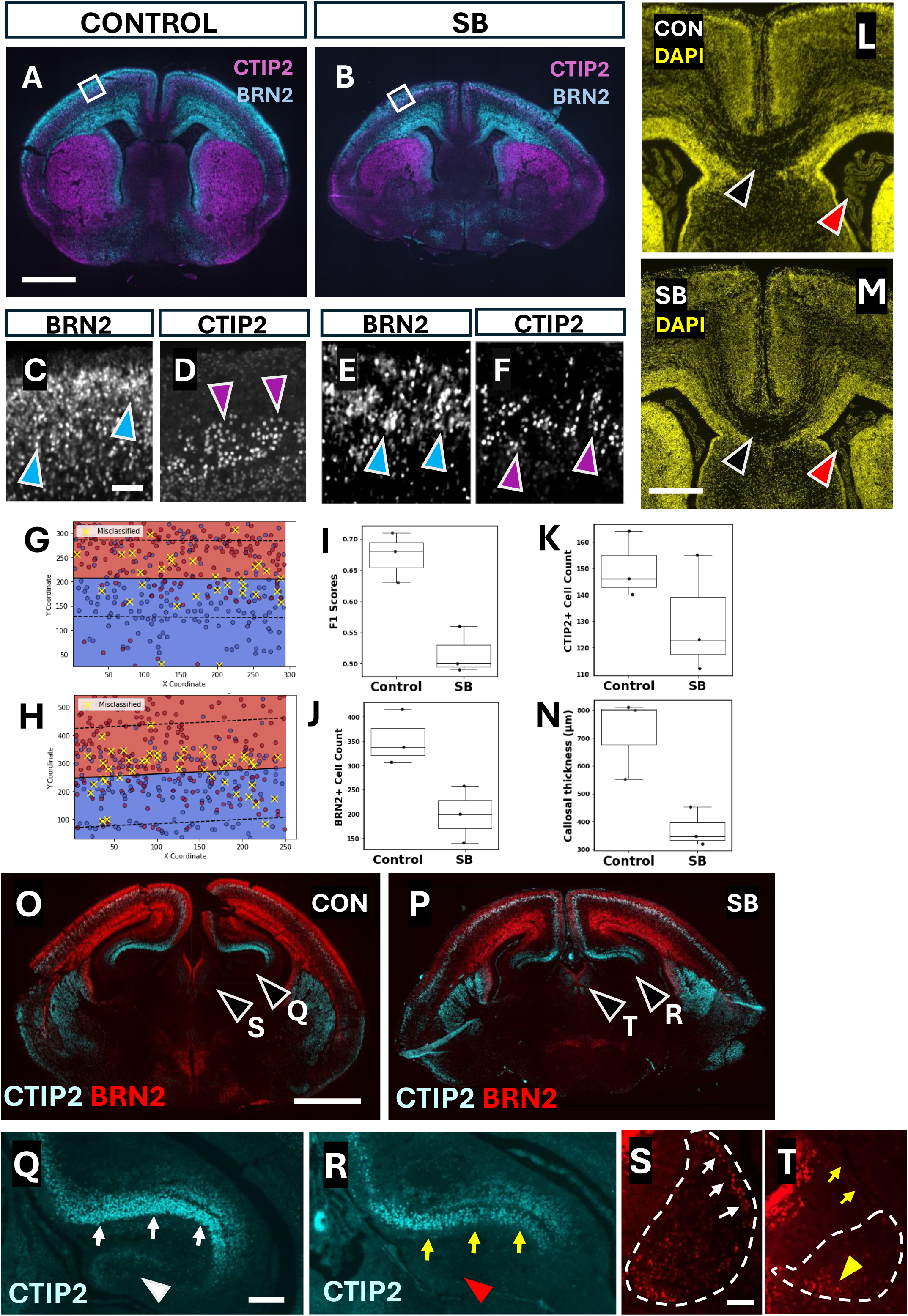
Neuronal heterotopia and agenesis in forebrain structures. (**A-F**) Representative coronal sections through the forebrain of control (A) and SB (B) at E18.5, immunostained with CTIP2 (magenta) and BRN2 (blue). Higher magnification (C-F) of boxed areas in (A, B) showing binarised BRN2+ (blue arrowheads) and CTIP2+ neurons (magenta arrowheads). There is greater mixing of CTIP2+ and BRN2+ neurons in SB than control. Scale bars: 1 mm in A, B; 0.05 mm in C-F). (**G, H**) Representative plots illustrating the decision-boundary learned by the linear Support Vector Machine (SVM) in control (G) and SB (H) cerebral cortex (see Materials & Methods). The boundary line, along with its parallel margins (dotted lines), is plotted over a mesh spanning the observed X-Y space. Each data point is coloured according to its original class label: BRN2 (red), CTIP2 (blue). Points highlighted in yellow are the misclassified instances. (**I**) F1 accuracy scores from the SVM analysis of control and SB (n = 3 for each) showing significantly lower scores in SB than control (Student’s t-test; p = 0.008), confirming that the two neuronal populations overlap more extensively in SB. (**J, K**) Comparison of BRN2+ (J) and CTIP2+ (K) cell counts in the cortex between control and SB (n = 3 for each). BRN2+ cells are significantly less abundant in SB than control (J; Student’s t-test; p = 0.03), whereas CTIP2+ cell number does not differ significantly (K; Student’s t-test; p = 0.25). (**L-N**) DAPI staining of coronal sections midway through the forebrain in control (L) and SB (M). Black arrowheads point to the corpus callosum, which appears thinner in SB. Red arrowheads point to choroid plexus of the lateral ventricles, which appears less convoluted and with a smaller area in SB. Scale bar: 0.5 mm in L, M. Quantification shows significantly reduced callosal thickness (N) in SB compared with control (Student’s t-test; p = 0.021). See Figure 4P for quantification of choroid plexus volumes, and Supplementary Figure 6 for additional corpus callosum comparison. (**O-T**) Coronal sections through control (O, Q, S) and SB (P, R, T) forebrain-midbrain regions, immunostained for CTIP2 (cyan) and BRN2 (red). Magnified views of hippocampus (Q, R) reveal lack of CTIP2+ cells in the dentate gyrus (compare white and red arrowheads in Q and R respectively) and fewer C2 and C3 cells (compare white and yellow arrows in Q and R respectively) in SB compared with control. Magnified habenula views (S, T) show a reduced overall area of BRN2+ cells (dashed lines), with few cells in a dorsolateral position (compare white and yellow arrows in S and T respectively) and an abnormally located cell cluster in the ventromedial region (yellow arrowhead in T). See also Supplementary Figures 7 and 8 for additional examples of hippocampus and habenula comparisons. Scale bars: 1 mm in O, P; 0.2 mm in Q, R; 0.05 mm in S, T.

### Callosal and hippocampal dysgenesis in SB brains

The corpus callosum and hippocampus are two brain regions reliant on timely neurogenesis, with known abnormalities in human Chiari II patients (Supplementary Table 1). As also detected at E15.5 (Figure 5K, L), callosal hypogenesis was found to be a feature of E18.5 SB brains, with significantly reduced callosal thickness compared to controls (Figure 7L-N; Supplementary Figure 6). At the same stage, we evaluated the presence of CTIP2+ and BRN2+ cells in the hippocampus and habenula (Figure 7O-T; Supplementary Figures 7,8). The region of the SB dentate gyrus entirely lacked CTIP2+ cells, and the CA1 and CA2 regions showed reduced cell numbers in SB compared with controls (Figure 7Q, R; Supplementary Figure 7). We also found a lack of BRN2+ cells at the lateral aspect of the habenular nuclei in SB, and a general heterotopic arrangement of cells ventro-medially (Figure 7S, T; Supplementary Figure 8). Together, these findings suggest the higher brain defects characteristic of Chiari II in humans are also present within our *Cdx^cre/+^;Pax3^fl/fl^* mouse model.

## DISCUSSION

Open SB, also called myelomeningocele, is a common birth defect in which the spinal cord fails to close during embryonic development. It affects on average 0.5 per 1000 live births worldwide, but with much higher frequencies in some resource-poor countries: e.g. 3-13 per 1000 in Ethiopia (Gebremariam et al., 2025). Fortification of the food supply with folic acid reduces SB prevalence by up to 50% (Atta et al., 2016), but at least a proportion of other cases are likely ‘folate-non-preventable’ (Greene et al., 2017). The increasing availability in many countries of fetal surgery for SB (Gallagher et al., 2023) offers an alternative to prenatal diagnosis and abortion, and may increasingly encourage parents to pursue postnatal survival of their SB-affected children. Hence, understanding and working towards improvement of the physical and mental health of children with SB is a research priority.

A focus in SB, postnatally, is often on lower body functional deficit (Oakeshott et al., 2012). However, many key long-term sequelae of SB result from brain anomalies, collectively termed the Chiari II malformation, which affects up to 90% of SB cases (Stevenson, 2004). This leads to hydrocephalus due to the associated hindbrain herniation, and also higher (‘supratentorial’) brain disorders, which include cortical heterotopias, callosal hypogenesis and basal ganglia anomalies (summarised in Supplementary Table 1). The higher brain defects are associated with frequent learning difficulties that can seriously affect the lives and independence of young people with SB (Rofail et al., 2013).

### Causation of Chiari II

The co-existence of Chiari II and SB has long indicated a significant link between the two conditions. However, it has been unclear whether Chiari II occurs secondary to an open spine or as a separate, primary defect that is related to SB (e.g. having a common genetic aetiology) but involving different direct effects of the causative factor. In our study, the *Cdx2^cre/+^;Pax3^fl/fl^* mouse maintains a genetically wild-type head despite a mutant lower body and SB phenotype. Nonetheless, we find typical Chiari II signs, including hindbrain herniation, posterior skull defects, global brain ‘collapse’ with posterior displacement, a number of forebrain defects including hippocampal and callosal agenesis, and heterotopia of the cortical layers. Given the multiple functions of Pax3 (Greene et al., 2009), it remains a formal possibility that an alteration other than open SB, in the Pax3-deleted low spine or surrounding tissues, led to the Chiari II phenotype we observed. Nevertheless, given the demonstrated association of Chiari II with SB in humans, where a variety of genetic and non-genetic causative factors have been implicated (Copp et al., 2013), our results provide strong experimental evidence that Chiari II can arise entirely secondary to SB. This places great importance on an improved understanding of the cause-and-effect relationship between SB and the origin of Chiari II.

### Evidence for disorders of brain neurogenesis and neuronal migration in Chiari II

We found cortical thinning and proliferative (ventricular) zone thickening in the telencephalon of the mouse fetuses with SB, raising the possibility of a brain neurogenesis disorder. Moreover, our finding of cortical heterotopias, in which neuronal sub-populations were intermixed in SB brains, as opposed to segregated in distinct layers, provides evidence for early-arising neuronal migration disorders. These results coincide with previous observations in humans: Fietz et al (2020) found a thickening of the subventricular zone (SVZ) by immunofluorescence of neuropathological specimens at 11-15 gestational weeks; Masse et al (2023) reported thickening of proliferative zones (VZ and SVZ) in human SB based on *in utero* MRI at 17-26 weeks; Juranek et al (2008) identified a series of brain morphometric changes in children aged around 12 years with open SB who had received CSF shunts for hydrocephalus and Chiari II. Hence, at varying stages of fetal and postnatal development, individuals with SB show higher brain anomalies, and this correlates with a series of specific neurocognitive disorders that can limit the education and independent living of children with SB (Rofail et al., 2013; Schneider et al., 2021). A challenge for future research is to determine the molecular and cellular pathways that are disturbed during brain neurogenesis in Chiari II, in order to arrive at potential preventive interventions.

### Origin of posterior skull disorders in Chiari II

A small, overcrowded posterior skull fossa is a recognised feature of Chiari II in humans (Juranek and Salman, 2010; Woitek et al., 2014), and this has been reproduced in chick embryos with surgically-created SB (Hwang et al., 2022; Sim et al., 2013). Interestingly, in the sheep model of surgically-induced SB, which provided the first demonstration of fetal surgical SB closure (Meuli et al., 1995), there is a lack of full-blown Chiari II disorder, and this has been attributed to the late creation of the artificial SB, long after the early events of brain and skull development are complete (Steele et al., 2020). McLone and Knepper (1989) proposed a small posterior fossa in Chiari II to result from failure of hindbrain expansion during development, and this idea fits well with accumulated research showing close alignment between brain and skull development (Richtsmeier and Flaherty, 2013). Our findings bear on this topic, as we identified different degrees of posterior bony skull defect in fetuses with SB. The most extreme situation involved a complete absence of the SO bone, and these fetuses showed a total lack of contact between hindbrain and developing posterior skull. Less severe SO reduction was associated with reduced hindbrain-skull contact. Hence, it seems possible that a sequence of developmental events, leading from brain expansion to posterior skull fossa induction, may be conserved in mice and humans and, when faulty, may account for the posterior skull phenotype in Chiari II.

### Role for CSF in Chiari II brain disorder

McLone and Knepper’s (1989) unified theory of Chiari II suggested CSF leakage through the SB lesion as the primary disruptor of embryonic brain development. Since that time, much circumstantial evidence has accumulated to support this idea. For example, Chiari II brain defects are found in open SB but not in skin-covered low spinal dysraphic conditions, where there is no CSF leakage (Schneider et al., 2021). Spinal cord ‘tethering’ occurs in both, hence ruling out downward traction as an alternative explanation for hindbrain herniation in Chiari II. Curtailment of CSF leakage by fetal surgery to close the SB lesion reduces hindbrain herniation in a proportion of cases, lowering the risk of hydrocephalus (Joyeux et al., 2018). However, the higher (supratentorial) defects in Chiari II persist into childhood, despite fetal surgery for SB (Calle et al., 2020), arguing for an earlier pathogenesis. Our finding that Chiari II brain defects occur in the mouse SB model, despite a wild-type head, provides the first experimental evidence for a secondary origin of Chiari II in relation to SB, which could be mediated through CSF leakage.

Embryonic CSF is rich in growth factors and signalling molecules that play major roles in guiding brain morphology, neural progenitor proliferation and CNS specification. For example, midbrain neuroepithelial explants from chick embryos fail to undergo neurogenesis, and show failed cell proliferation and apoptosis, in the absence of CSF (Gato et al., 2005). Moreover, CSF of embryonic origin supports neuronal differentiation from cultured stem cells (Shokohi et al., 2018), and stimulates more neurogenesis than adult CSF in embryonic brain slices (Alonso et al., 2017). Adult CSF promotes glial (astrocytic) rather than neuronal differentiation (Buddensiek et al., 2009).

Prior to neural tube closure, amniotic fluid bathes the open neuroepithelium, including the future brain, whereas CSF composition diverges from amniotic fluid soon after the neural tube has closed and sealed (Chau et al., 2015). Identified CSF components include serotonin, retinoids, Shh, LIF, Fgf2, Igf2, Wnts, nanovesicles and exosomes: all involved in neural progenitor survival, differentiation and brain morphogenesis (Fame and Lehtinen, 2020). In the case of SB, CSF composition was found to differ between unaffected children and those with postnatally-repaired SB (Pal et al., 2005). Since CSF leaks continuously from the open spinal lesion, this may reduce pressure within the cranium, but also could alter the availability of molecules essential for neurogenesis. Hence, there is a strong precedent for CSF compositional change affecting neuronal development, with a key role in the neurogenesis disorders of Chiari II.

## Conclusion

We have developed a mouse genetic model which demonstrates that Chiari II can arise secondary to SB, rather than as a separate primary lesion. Given the close similarity between mouse and human SB, this suggests that human Chiari II may also be entirely secondary to an open SB. We show that the mouse model exhibits a wide range of brain anomalies, typical of human Chiari II, and further present evidence that these may largely arise from early disorders of brain neurogenesis and neuronal migration. These fundamental events of brain development are largely complete by the time that fetal surgery is performed for SB, and so cannot be reversed, unlike hindbrain herniation. Our work will now focus attention on the causative SB-to-Chiari II link which may in future become susceptible to clinical interventions to prevent Chiari II defects, even in the presence of SB.

## MATERIALS AND METHODS

### Mouse procedures

Mouse research was reviewed and approved by the Animal Welfare and Ethical Review Body of University College London, and authorised by a Project Licence (PP0411055) under the auspices of the UK Animals (Scientific Procedures) Act 1986. Pregnant female mice served as a source of embryos and fetuses (of both sexes) following mating with stud males. *Cdx2^cre/+^*male mice (Hinoi et al., 2007) were obtained from the Jackson Laboratory (strain 009350: B6.Cg-Tg(CDX2-cre)101Erf/J) and maintained by breeding with C57BL/6J females. Genotyping was by PCR for Cre genomic sequence (Danielian et al., 1998). *Pax3^fl/+^* mice were a gift of Dr Simon Conway and kindly provided by the Pasteur Institute, Paris. Inter-crosses produced *Pax3^fl/fl^* offspring which were maintained as a homozygous colony. Genotyping was as described (Koushik et al., 2002).

Experimental litters were produced by mating *Cdx2^cre/+^;Pax3^fl/+^*males with *Pax3^fl/fl^* females (Figure 1A), to produce 4 offspring genotype categories (Table 1). *Cdx2^cre/+^;Pax3^fl/fl^* embryos and fetuses provided the model of SB/Chiari II, while non-Cre littermates (*Pax3^fl/+^*or *Pax3^fl/fl^*) served as controls. Reporter mTmG mice (JAX strain 007576; *Gt(ROSA)26Sor^tm4(ACTB-tdTomato,-EGFP)Luo^*/J)(Muzumdar et al., 2007) and ROSA26^eYFP^ (JAX strain 006148; B6.129X1-*Gt(ROSA)26Sor^tm1(EYFP)Cos^*/J) females were bred with *Cdx2^cre/+^* males to generate embryos for *Cdx2^cre^* recombination pattern analysis.

Pregnant dams were culled by cervical dislocation, with death confirmed via cutting the femoral artery. Embryos and fetuses (E10.5-E18.5) were dissected from the uterus in Dulbecco’s Modified Eagle’s Medium (DMEM) containing 10% fetal bovine serum, rinsed in phosphate buffered saline (PBS) and fixed in 4% paraformaldehyde in PBS (PFA) at 4°C overnight for histological analysis or in 4% PFA for 7 days then 1% PFA for microCT scanning. Yolk sacs of individual embryos and fetuses were rinsed in PBS and stored for genotyping. Owing to the perinatal lethality of SB in mice, it was not possible to study postnatal features of Chiari II in the mouse model.

### Cryosection preparation

Embryos and fetuses were fixed by immersion in 4% PFA for varying periods (overnight to several days) depending on stage, washed in PBS, dehydrated in increments of 10%, 20% and 30% sucrose in PBS, and kept at 4°C overnight or until the sample sank to the bottom of the tube. Embryos and fetuses were dissected further (brain only for E18.5 coronal sections; entire body for sagittal sections; head only for E15.5 coronal sections), patted dry from sucrose and placed in OCT (Optimal Cutting Temperature; SciGen, Oxford Instruments: 51-1625-0019). Once manoeuvred to desired cutting positions in embedding moulds, samples were frozen on dry ice and stored at -80°C. Cryosections were prepared using a Bright cryostat at 15 μm thickness, mounted on Superfrost Plus™ slides (Fisher Scientific: 10149870) and stored at -80°C until stained.

### Haematoxylin and eosin (H&E) staining

Slides were washed in double distilled (dd) H_2_O, then stained in filtered Harris’s haematoxylin (Sigma Aldrich: HH516) for 4 min. Slides were further rinsed in H_2_O and differentiated in acid/alcohol (1% HCl in 70% ethanol), rinsed in ultrapure (UP) H_2_O and ‘blued up’ in saturated lithium carbonate. Following another UPH_2_O wash, slides were stained with filtered aqueous Eosin Y (Sigma Aldrich: HT110232), washed in UPH_2_O, dehydrated in 100% ethanol, cleared in Histoclear (National Diagnostics; Scientific Laboratory Supplies: NAT1330) and coverslip-mounted using DPX (Sigma-Aldrich: 06522).

### Immunohistochemistry

Primary antibodies (all used at 1:200 dilution) were: anti-BRN2, rabbit polyclonal (Cell Signaling Technology: 12137S); anti-CASP (CUX1), mouse monoclonal IgG1 (ABCAM: ab54583); anti-CTIP2, rat monoclonal (ABCAM: ab184650); anti-PAX3, mouse monoclonal (Developmental Studies Hybridoma Bank: AB_528426); anti-TBR1, rabbit polyclonal (ABCAM: ab31940). Secondary antibodies were: goat anti-mouse IgG1 Alexa Fluor™ 568 (Invitrogen; A21124), donkey anti-rabbit Alexa Fluor™ 488 (Invitrogen; A32790) and chicken anti-rat Alexa Fluor™ 647 (Invitrogen; A21472). DAPI (4’,6-diamidino-2-phenylindole) nuclear dye was used at 1:5000 (ThermoFisher Scientific).

Cryosectioned tissue was processed for antigen retrieval using sodium citrate (0.1 M, pH 6.0) and warmed to 120°C in a decloaking chamber for 45 min. Slides were removed from the chamber and 2 ml PBS was applied for 10 min at room temperature (RT). PBS was removed and replaced by blocking buffer (10 ml PBS, 100 μL heat inactivated sheep serum, 100 μL 10% Triton-x100) for 1 h at RT. Primary antibody in blocking buffer was then applied, overnight at 4°C. Slides were washed 3x in PBS for 10 min each on a slow speed shaker. PBS was removed and secondary antibody in blocking buffer was added to the slides and left for 1 h at RT. DAPI was added to the slides for 10 min, followed by washes 3x in PBS. Sudan black (in 70% ETOH) was added for 10 min to minimise autofluorescence. A further 3-5 PBS washes were used if necessary to remove excess staining. Slides were mounted in Prolong Gold mounting medium (Thermo Fisher Scientific: P36934) beneath coverslips. Controls for immunostaining were routinely performed, including omission of primary antibody and staining of genetically null tissue, which generated an absent signal in the anti-Pax3 experiments.

### MicroCT

MicroCT preparation and scanning was carried out in the Embryo Phenotyping facility of the Mary Lyon Centre at MRC Harwell, Oxford, UK.

#### Hard tissue scanning

Fetuses were dissected in ice cold phosphate buffered saline (PBS) and exsanguinated by severing the umbilical vessels. After washing in PBS, fetuses were fixed by immersion in 4% PFA for 7 days at 4°C on a rocker, before storage in 1% PFA at 4°C. Samples were rinsed through 3 changes of ddH_2_O before embedding within an acrylic mount in 1% agarose, which was allowed to set for at least 1 h. MicroCT data sets were acquired using a Skyscan 1272 scanner (Bruker) with the x-ray source set to 70 kV and using 0.5 mm aluminium filter. Pixel resolution of 15 µm/pixel was set at 3x3 camera pixel binning (1344 x 896). Four projections were acquired and averaged every 0.4° through a total rotation of 360°, with random movements set to 25. NRecon (Bruker) was using for 3D reconstruction to PNG.

#### Soft tissue scanning

After storage in 1% PFA at 4°C prior to scanning, fetuses were contrasted by immersion in 50% Lugol’s solution (Sigma-Aldrich: 32922. 1:1 Lugol:ddH_2_O) at room temperature for 2 weeks on a rocker, protected from light, with the solution replaced every 2 days. Samples were then rinsed and incubated in ddH_2_O for at least 1 h before embedding within an acrylic mount in 1% agarose which was allowed to set for at least 2 h. MicroCT data sets were acquired using a Skyscan 1272 scanner (Bruker) with the X-ray source set to 80 kV and using 1 mm aluminium filter. Pixel resolution of 12 µm/pixel was set at 2x2 camera pixel binning (2048 x 2048). Four projections were acquired and averaged every 0.15° through a total rotation of 360°, with random movements set to 25. NRecon (Bruker) was using for 3D reconstruction to PNG using a constrained volume of 1000 x1400 pixels (width x height).

### Support vector machine learning analysis

Python 3.11.3 and its pandas, matplotlib and scikit-learn packages were used. For each sample, two datasets were combined: BRN2+ and CTIP2+ cell coordinates, as 2-dimensional X and Y values. These were labelled as 0 (BRN2+) and 1 (CTIP2+). After merging, the final dataset consisted of all measurements from both conditions. This dataset was then split into training (70%) and test (30%) subsets using a random state of 42 to ensure reproducibility. A linear Support Vector Machine (SVM) was trained on the ‘X‘,‘Y‘‘X’, ‘Y’‘X‘,‘Y‘ features in the training set to distinguish between the two labelled groups. Following training, the model’s performance was evaluated by generating predictions on the unseen test set. A confusion matrix revealed how many points were correctly classified in each category versus those that were misclassified. A classification report provided an F1 score of the model’s accuracy, while a harmonic mean of precision and recall of the SVM indicated its overall performance. Script for this analysis is available upon request.

### Statistical analysis

Statistical tests and graph preparation were carried out using Sigmaplot 14.5 and 16, and Python 3.11.3 and the pandas, matplotlib, seaborn, scipy, and numpy packages. Python scripts are available upon request.

## Acknowledgements

We thank Rosie Bunton-Stasyshyn, Jacqueline Horn and James Cleak of the Mary Lyon Centre at MRC Harwell for expert assistance with microCT.

## Author contributions

Conceptualization: M.C., A.J.C.; Data curation: M.C., A.J.C.; Formal analysis: M.C., A.J.C.; Funding acquisition: N.D.E.G., A.J.C.; Investigation: M.C., T.J.E., D.S., N.S.; Methodology: M.C., T.J.E., D.S., E.P.; Supervision: E.P., A.J.C.; Validation: G.L.G., N.K., A.J.C.; Writing – original draft: M.C., A.J.C.; Writing – review & editing: all authors.

## Data and resource availability

The data that support the findings of this study are available from the corresponding author, upon reasonable request.

---

**Supplementary Figure 1.**
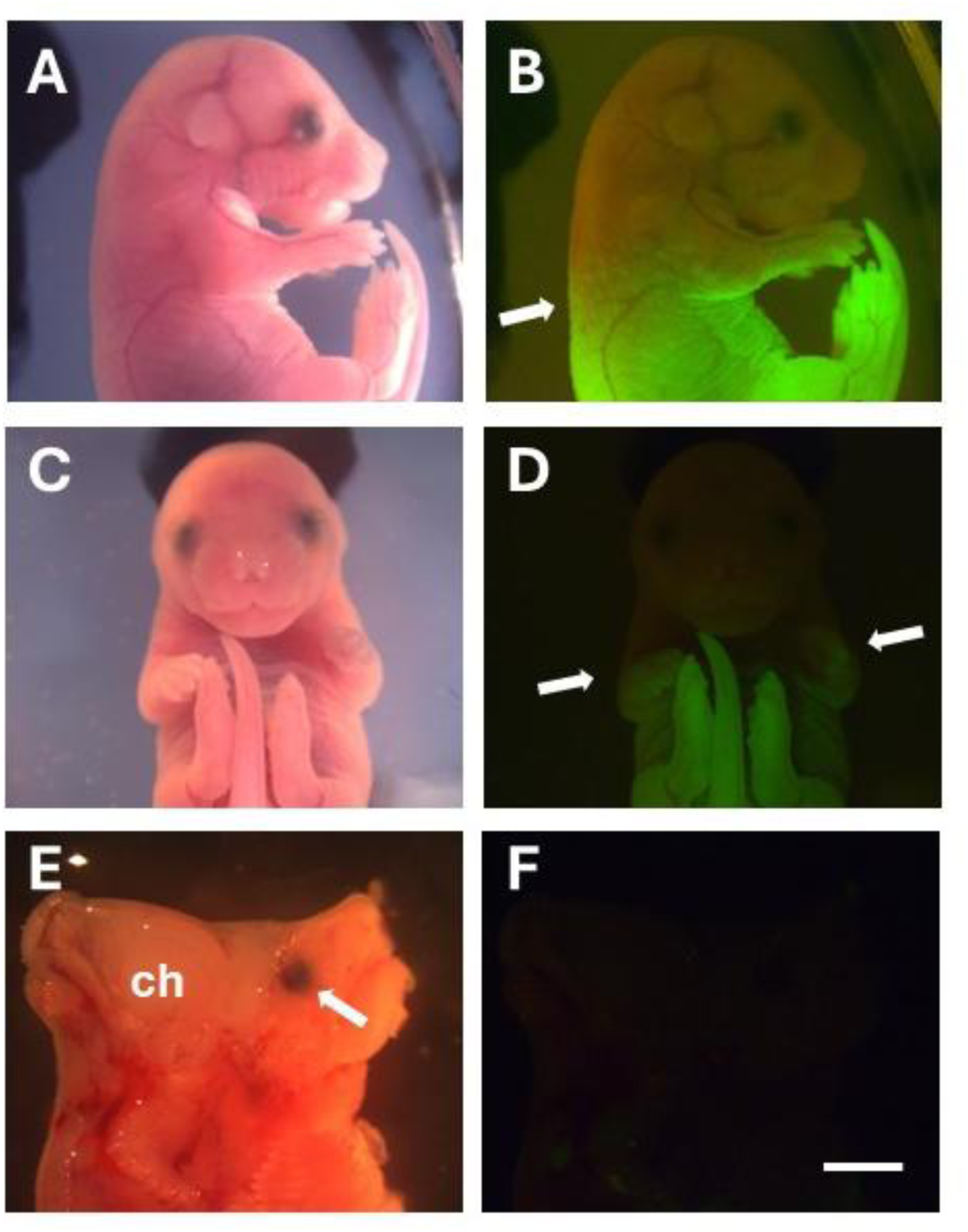
Analysis of rostro-caudal level of *Cdx2^cre^* recombination in E18.5 fetuses. (A-D) Bright-field (A,C) and fluorescence (B,D) images of E18.5 *Cdx2^cre/+^; Rosa26^eYFP^* fetuses, viewed from the right side (A,B) and front (C,D). The cut-off level of *Cdx2^cre^*-mediated eYFP recombination (arrows in B.D) is at the forelimb level, which corresponds to the thoraco-cervical boundary. (E,F) Fetal head bisected sagittally from the front (E; arrow indicates left eye; ch: cerebral hemisphere) to identify any evidence of brain Cdx2^Cre^ recombination, but none is seen in the fluorescence image (F). This analysis confirms our findings at E10.5 and E15.5 (Figures 1B,C; 2A-D) that *Cdx2^cre^* mediates recombination in the trunk region only, and not in the head. Sample size = 3. Scale bar = 2 mm in F (for A-F).

**Supplementary Figure 2.**
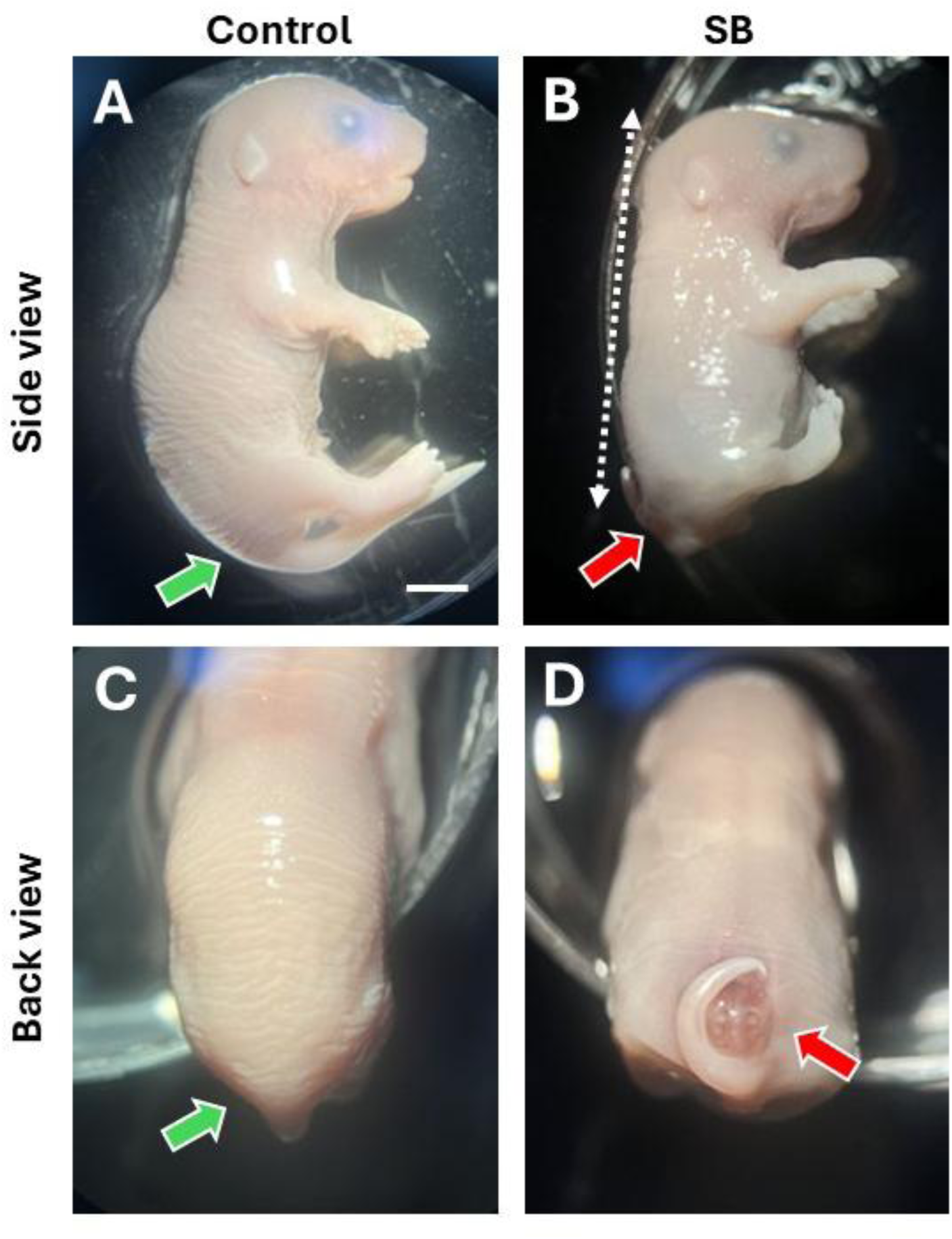
Additional views of E18.5 control and SB fetuses. (A,C) Control and (B,D) SB fetuses shown from the right side (A,B) and back (C,D) views. Green arrows: normal low spinal region in control. Red arrows: open SB with a curled tail defect. Note the abnormally short body axis in the SB fetus (dotted arrow). Scale bar = 2 mm in all parts.

**Supplementary Figure 3.**
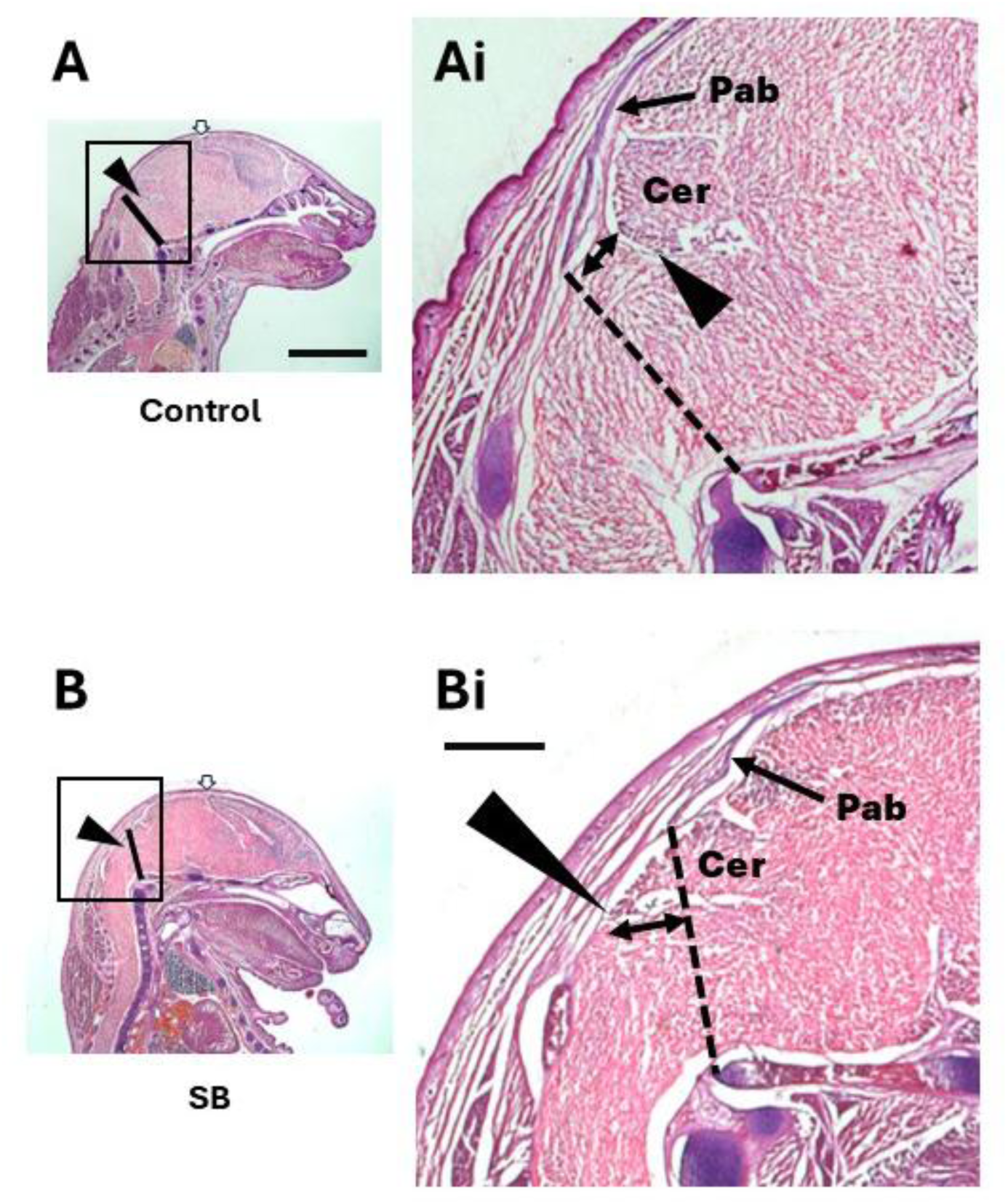
Analysis of hindbrain herniation in SB fetuses, compared with controls at E18.5. Higher magnification views (Ai, Bi) of the H&E-stained sagittal head sections (A, B) as reproduced from Figure 3A, B. A line was drawn from the caudal edge of the parietal bone dorsally (Pab in Ai, Bi) to the inflection point of the skull base ventrally, where it joins the vertebral column (purple). This line (dashed in Ai, Bi) is taken as the boundary between skull and spinal region. In the SB head, the caudal edge of the cerebellum (Cer; black arrowhead in Bi) is located below the line (double headed arrow in Bi), indicating a herniation. In contrast, the control head shows the cerebellum entirely above/rostral to the line (double headed arrow in Ai) indicating no herniation. See Figure 3C, D for quantitation of skull boundary and cerebellar herniation. Parietal bone length was measured from its rostral edge (open arrows in A, B) to its caudal edge (Ai, Bi), and normalised to antero-posterior head length in each fetus. Normalised parietal bone length does not differ between control and SB (Student’s t-test: p = 0.372; n = 3 for each genotype). Scale bars = 2 mm in A, B; 0.5 mm in Ai, Bi.

**Supplementary Figure 4.**
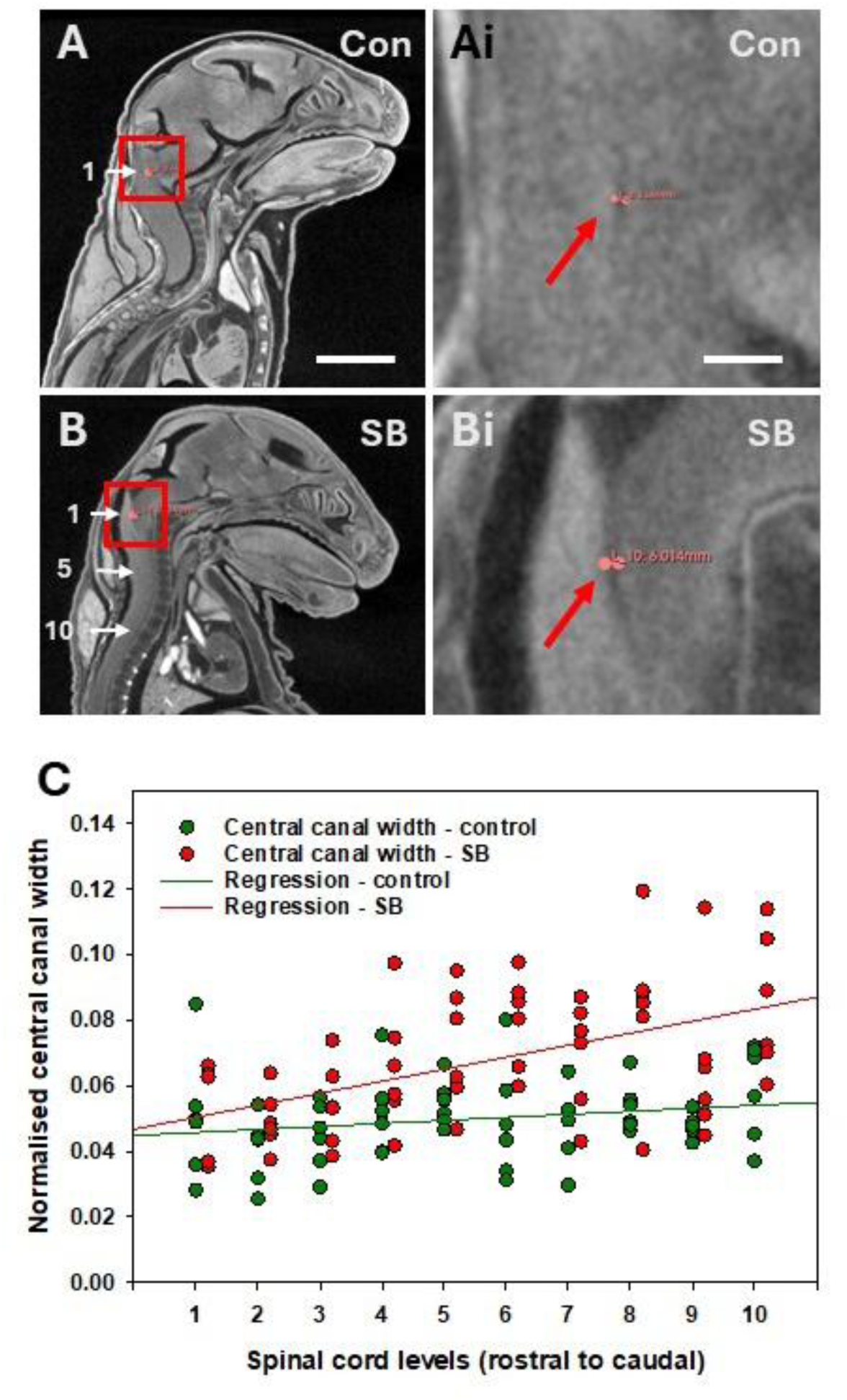
Enlargement of the central spinal canal in SB fetuses. (A-Bi) Soft-tissue microCT images of control (Con; A, Ai) and SB (B, Bi) E18.5 fetal heads at low magnification (A, B) and with magnified views of the boxed areas (Ai, Bi). The central canal is visible as a dark vertical line against the grey spinal cord tissue in Ai, Bi (red arrows). Central canal width could be reproducibly measured in these sagittal images, and was determined at ten equally spaced levels, 1-10, moving down the spinal cord (numbered white arrows in A, B). Scale bars = 2 mm in A, B; 0.2 mm in Ai, Bi. (C) Central canal width, normalised to spinal cord dorso-ventral width, plotted against spinal cord level in control and SB fetuses (n = 6 each). Statistical analysis: 2-way ANOVA shows significant differences between genotype (control vs SB) and spinal cord level (p < 0.001 for each). Linear regression equations: Y = 0.0009 X + 0.04483 (control) and Y = 0.0037 X + 0.0467 (SB). R^2^ = 0.30 (control) and 0.71 (SB). Student t-tests show significant differences between regression line slopes (p = 0.011) but not Y-axis intercepts (p = 0.762).

**Supplementary Figure 5.**
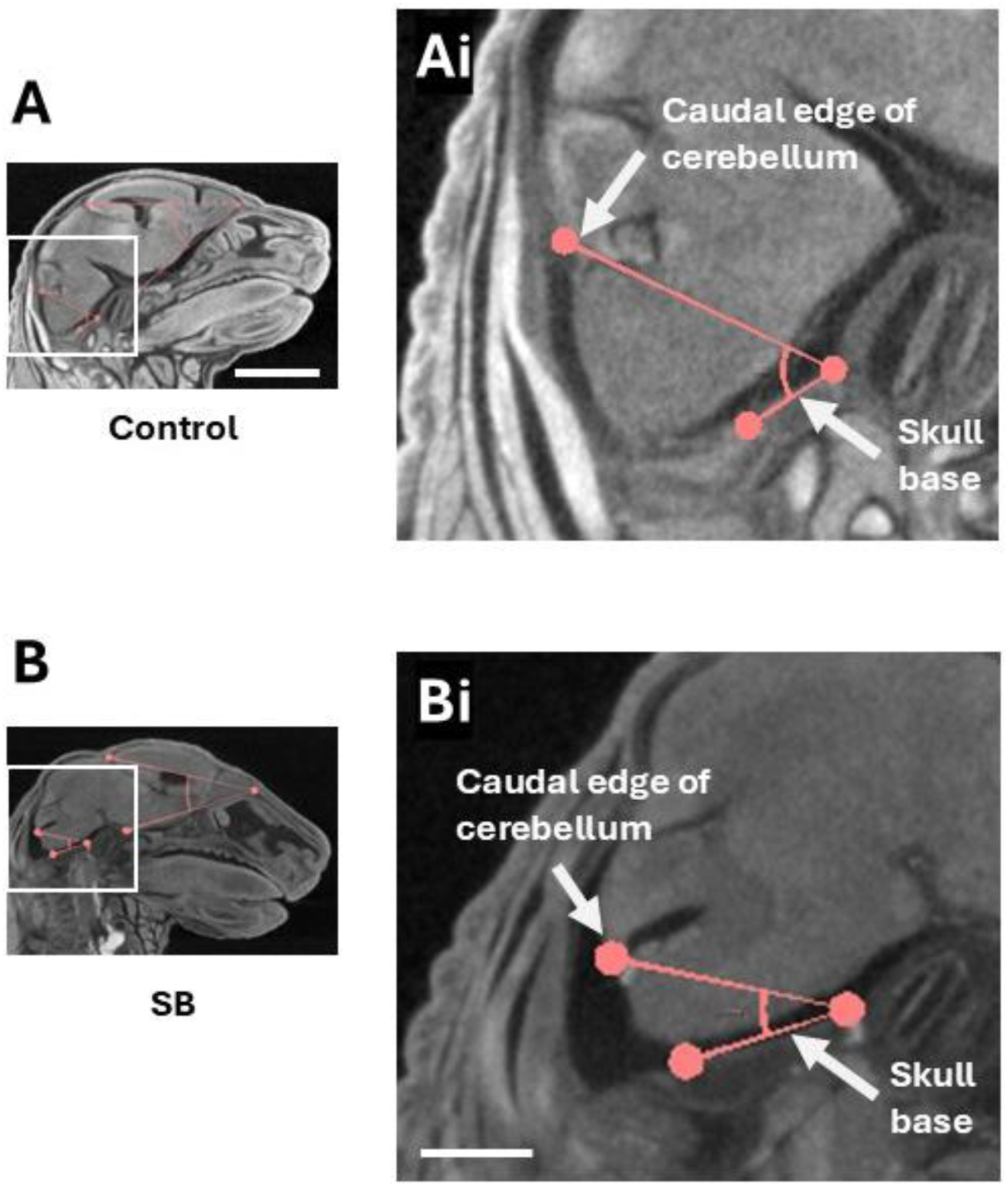
Analysis of cerebellar position in relation to the skull base of SB and control fetuses at E18.5. Higher magnification views (Ai, Bi) are shown of the soft tissue microCT images (A, B) as reproduced from Figure 4J, K. Angle x (caudal edge of cerebellum to inflection point of the skull base) is smaller in the SB head (Bi) than in control (Ai), consistent with hindbrain herniation in the SB fetus. See Figure 4 N for quantitation. Scale bars = 2 mm in A, B; 0.5 mm in Ai, Bi.

**Supplementary Figure 6.**
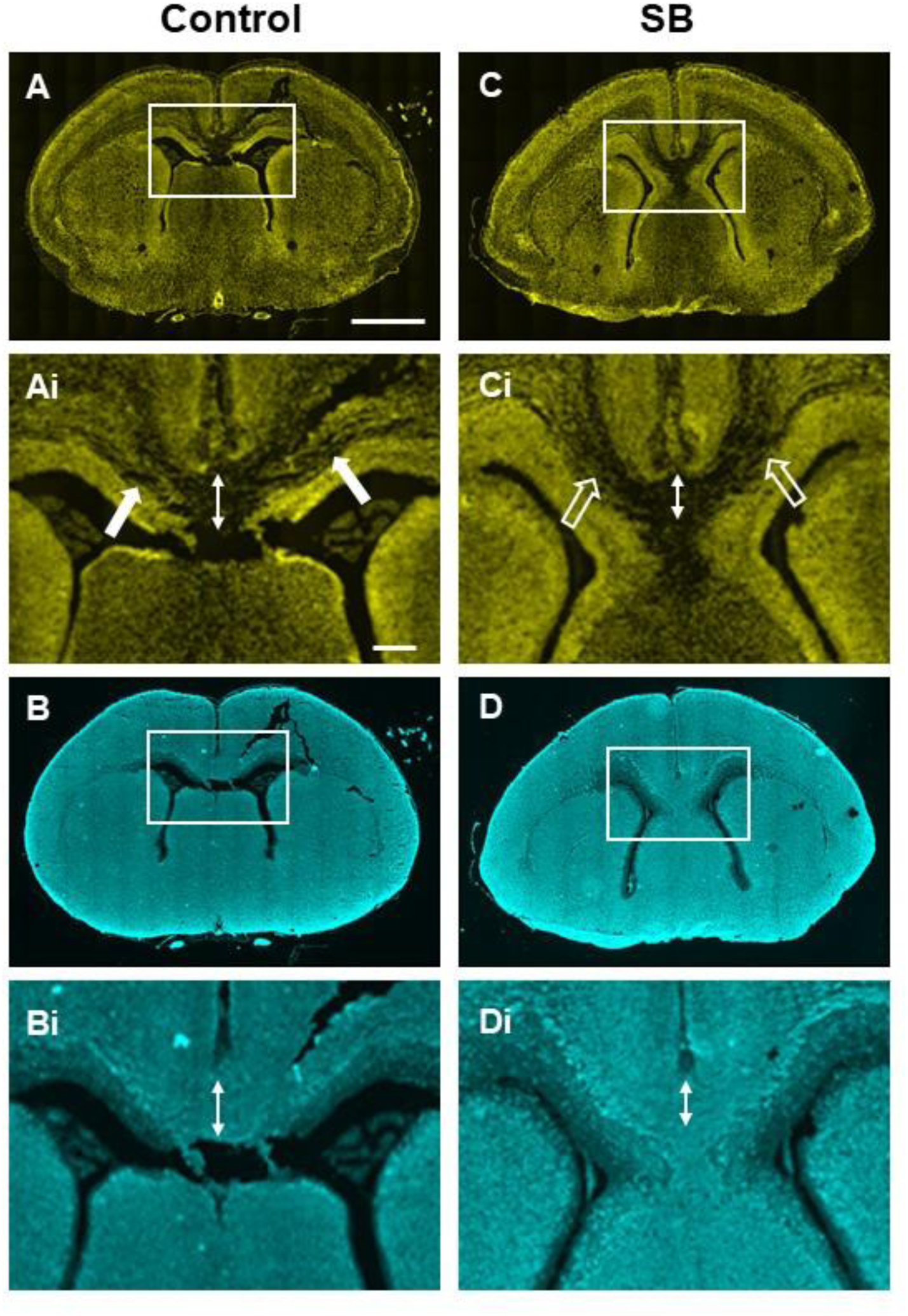
Further example of callosal hypogenesis in an SB fetus at E18.5. Coronal sections through the brains of control (A, B) and SB (C, D) fetuses at low magnification, with boxed areas shown at higher magnification (Ai-Di). Sections stained with DAPI (A, Ai, C, Ci) and by Tuj1 immunohistochemistry (B, Bi, D, Di). Double headed arrows (Ai-Di) indicate the dorso-ventral thickness of the corpus callosum, which is reduced in the SB fetus (Bi, Di) compared with control (Ai, Ci). Note the abundant crossing fibres visible by DAPI staining that are entering the midline of the control fetus (solid arrows in Ai), which are greatly reduced in the SB fetus (open arrows in Bi). Scale bars = 1 mm in A -D; 0.2 mm in Ai-Di.

**Supplementary Figure 7.**
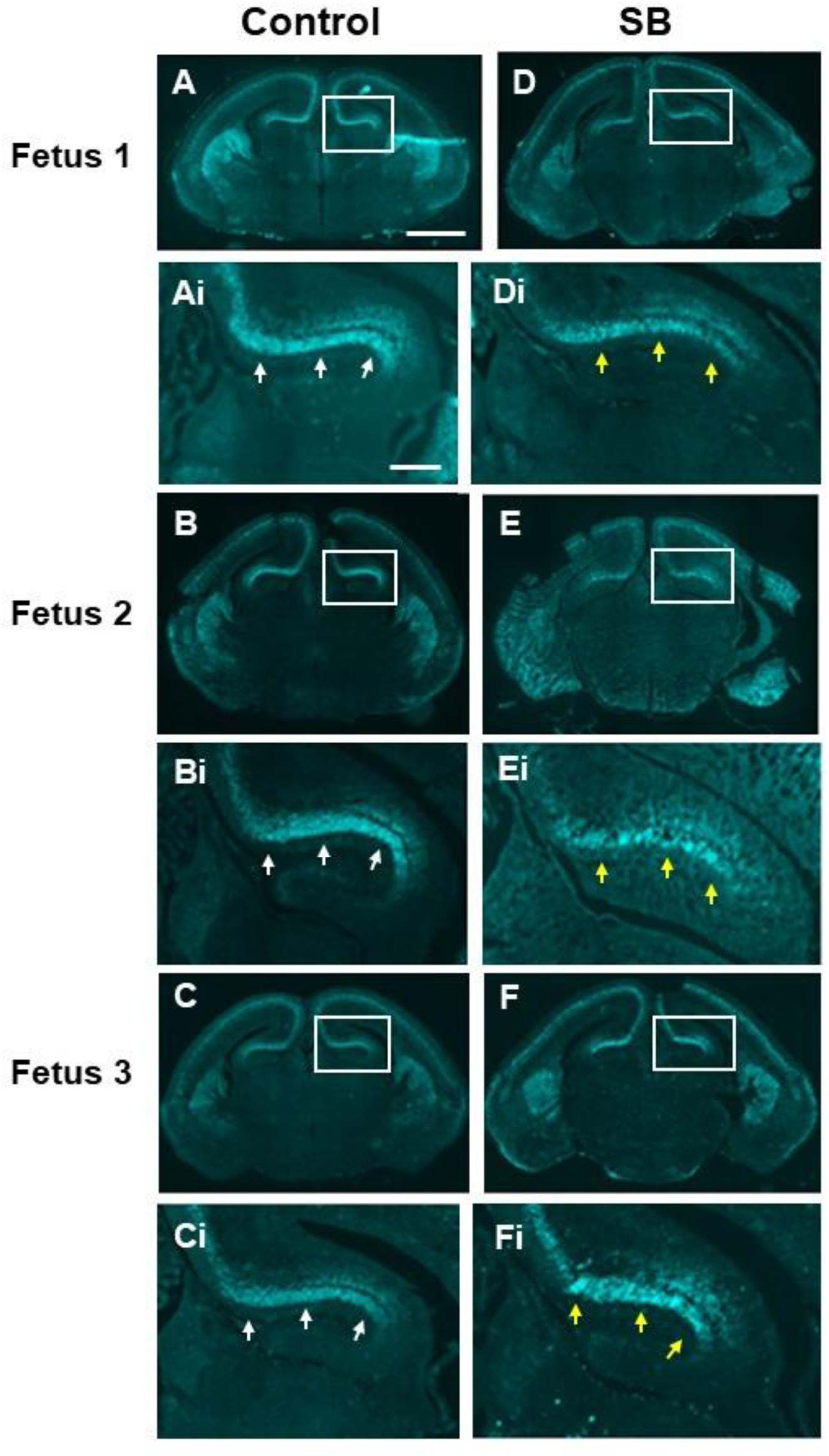
Further examples of hippocampal hypogenesis in SB fetuses at E18.5. Coronal sections through the brains of three separate control (A-C) and SB (D-F) fetuses at low magnification, with boxed areas shown at higher magnification (Ai-Fi). Sections stained by CTIP2 immunohistochemistry. The CA1/CA2 regions of all three SB fetuses show reduced cellularity (yellow arrows in Di-Fi) compared with controls (white arrows in Ai-Ci). Control fetus 2 and SB fetus 1 are also shown (in adjacent sections) in Figure 7Q,R. Scale bars: 1 mm in A-F; 0.2 mm in Ai-Fi.

**Supplementary Figure 8.**
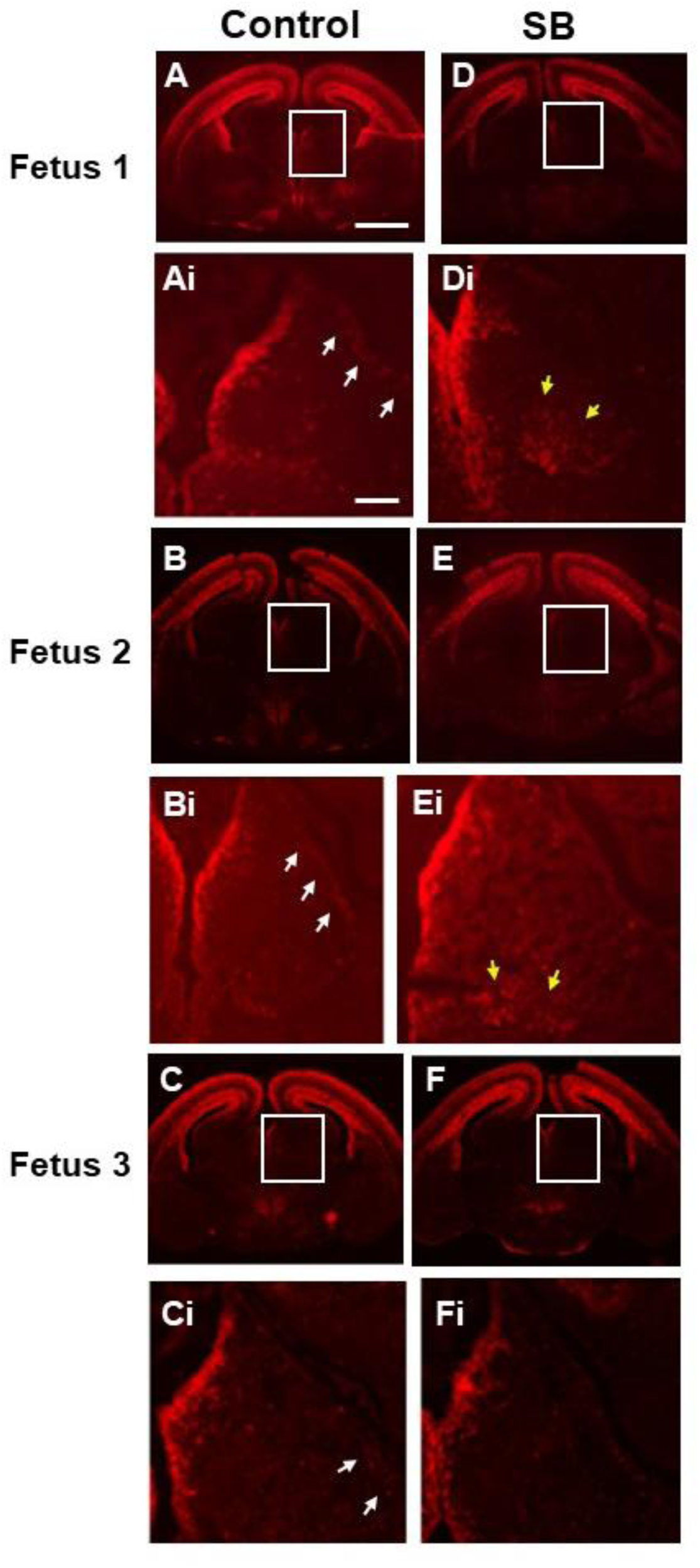
Further examples of habenula hypogenesis in SB fetuses at E18.5. Coronal sections through the brains of three separate control (A-C) and SB (D-F) fetuses at low magnification, with boxed areas shown at higher magnification (Ai-Fi). Sections stained by BRN2 immunohistochemistry. White arrows indicate a slanting line of BRN2+ cells in the lateral habenula of two control brains (Ai, Bi), with a few cells visible at this location in the third control (Ci) brain. In contrast, such lateral habenular BRN2+ cells are not visible in any of the SB fetuses, although two show apparently increased BRN2+ cellularity in the ventromedial habenular region (yellow arrows in Di and Ei). Scale bars: 1 mm in A -F; 0.2 mm in Ai-Fi.

**Supplementary Table 1.** Brain anomalies in human Chiari II.

| Brain region | Chiari II description |
| --- | --- |
| Cerebellar vermis & 4<sup>th</sup> ventricle | Downward displacement, peg-like shape <sup>1</sup> |
| Cisterna magna | Absent, producing “banana sign” <sup>2</sup> |
| Cerebellum | Folia shallow or absent <sup>3</sup>; heterotopia <sup>4</sup> |
| Tentorium cerebelli | Dysplastic <sup>5</sup>; low lying <sup>6</sup> |
| Cortex | Reduced thickness, heterotopia, smaller occipital surface areas <sup>7, 8</sup> |
| White matter | Total significantly reduced <sup>8</sup> |
| Ependyma | Denudation and rosettes <sup>4</sup> |
| Global neuronal nuclei | Varied heterotopia only present in SBA not in SBO <sup>4</sup> |
| Corpus callosum | Anomalies/partial agenesis <sup>4, 9</sup>, hypoplasia (splenium and truncus affected most frequently) <sup>10</sup> |
| Pons | Hypoplastic <sup>10</sup> |
| Meninges | Dysplasia of subarachnoid space <sup>4</sup> |
| Ventricular system | Hydrocephalus, aqueductal stenosis <sup>4</sup>; ventriculomegaly <sup>11</sup>; enlarged lateral ventricles prenatally <sup>9</sup> |
| Mesencephalon | Hypoplastic <sup>10</sup> |
| Gyri | Stenogyria <sup>4, 10, 12</sup>; polymicrogyria <sup>12</sup> |
| Medulla | Medullary kinking <sup>10</sup> |
| Tectum | Tectal beaking <sup>5, 11</sup> |
| Colliculus | Collicular fusion <sup>11</sup> |
| Massa intermedia | Enlargement <sup>13</sup> |
| Habenular commissure | Elongation <sup>13</sup> |
| Pineal gland | Elongation <sup>13</sup> |
| Periventricular zone | Nodular heterotopia <sup>14</sup> |
| Cranial nerves | Hypoplasia <sup>15</sup> |
| Fetal proliferative zones (ventricular & subventricular) | Larger volumes <sup>9</sup> |
| Diencephalon (fetal) | Decreased volume <sup>9</sup> |

